# Long read sequencing of 3,622 Icelanders provides insight into the role of structural variants in human diseases and other traits

**DOI:** 10.1101/848366

**Authors:** Doruk Beyter, Helga Ingimundardottir, Asmundur Oddsson, Hannes P. Eggertsson, Eythor Bjornsson, Hakon Jonsson, Bjarni A. Atlason, Snaedis Kristmundsdottir, Svenja Mehringer, Marteinn T. Hardarson, Sigurjon A. Gudjonsson, Droplaug N. Magnusdottir, Aslaug Jonasdottir, Adalbjorg Jonasdottir, Ragnar P. Kristjansson, Sverrir T. Sverrisson, Guillaume Holley, Gunnar Palsson, Olafur A. Stefansson, Gudmundur Eyjolfsson, Isleifur Olafsson, Olof Sigurdardottir, Bjarni Torfason, Gisli Masson, Agnar Helgason, Unnur Thorsteinsdottir, Hilma Holm, Daniel F. Gudbjartsson, Patrick Sulem, Olafur T. Magnusson, Bjarni V. Halldorsson, Kari Stefansson

## Abstract

Long-read sequencing (LRS) promises to improve characterization of structural variants (SVs), a major source of genetic diversity. We generated LRS data on 3,622 Icelanders using Oxford Nanopore Technologies, and identified a median of 22,636 SVs per individual (a median of 13,353 insertions and 9,474 deletions), spanning a median of 10 Mb per haploid genome. We discovered a set of 133,886 reliably genotyped SV alleles and imputed them into 166,281 individuals to explore their effects on diseases and other traits. We discovered an association with a rare (AF = 0.037%) deletion of the first exon of *PCSK9*. Carriers of this deletion have 0.93 mmol/L (1.31 SD) lower LDL cholesterol levels than the population average (p-value = 7.0·10^−20^). We also discovered an association with a multi-allelic SV inside a large repeat region, contained within single long reads, in an exon of *ACAN*. Within this repeat region we found 11 alleles that differ in the number of a 57 bp-motif repeat, and observed a linear relationship (0.016 SD per motif inserted, p = 6.2·10^−18^) between the number of repeats carried and height. These results show that SVs can be accurately characterized at population scale using long read sequence data in a genome-wide non-targeted approach and demonstrate how SVs impact phenotypes.

---

Human sequence diversity is partially due to structural variants^1^ (SVs); genomic rearrangements affecting at least 50 bp of sequence in forms of insertions, deletions, inversions, or translocations. The number of SVs carried by each individual is less than the number of single nucleotide polymorphisms (SNPs) and short (< 50 bp) insertions and deletions (indels), but their greater size makes them more likely to have a functional role^2^, as evident by their disproportionately large impact on diseases and other traits^2,3^.

Extensive characterization of three trios sequenced using several technologies^4^ and an annotated set based on one sample (HG002)^5^ indicate that humans carry 23-31 thousand SVs per individual. Most studies using whole genome sequence data are based on short read sequencing (SRS), where reads are typically 100-200 bp in length, allowing SNPs and small indels to be reliably identified^6,7^. However, short reads make the discovery, genotyping and characterization of SVs difficult^8^ and the number of SVs found per individual has been limited to 2-11 thousand in large scale studies using SRS^3,9–12^. Long-read sequencing (LRS), with read lengths of several kilobases (kb), allows SVs to be detected with greater accuracy. The typical process for identifying SVs involves mapping and comparing sequence reads to a reference genome. Due to their greater length, LRS reads can be mapped more accurately than SRS reads^8^. LRS reads are also more likely to cover entire SVs, enabling better determination of their breakpoints and length. However, LRS reads have a relatively high sequencing error rate (often more than 10%), that varies depending on sample quality, sequencing technology, and protocol^8^. High error rates can result in artifacts^8,13^, as well as failure in SV identification. Artifacts can be especially challenging in large scale studies, where accumulating false-positives may dominate results and hinder downstream analysis, such as genome-wide associations. Although there exist studies on detecting and characterizing SVs in human genomes using long-reads^8,14–17^ on select small datasets, analysis at scale has not been reported.

We present the first application of LRS at a population scale, focused on identifying a set of reliable SVs consistently called across individuals which can be used for downstream analysis in the context of diseases and other traits. We sequenced 3,622 Icelanders using Oxford Nanopore Technologies (ONT), including 441 parent-offspring trios, recruited for various studies at deCODE genetics^18^. DNA was isolated from whole blood (N = 3,524) and heart tissue (N = 102) and sequenced with ONT PromethION instruments (Methods). SRS and DNA chip data were also available for all of these individuals^19^. We introduced a number of tools and approaches to facilitate SV analysis using long reads characterized by a high error rate at scale, including SV filters, and heuristics for merging SVs. Finally, to illustrate the power of population based LRS, we developed a tool to perform joint genotyping on LRS data.

We basecalled raw sequence data in 4,757 flow cells, where half of all sequenced basepairs (N50) belonged to reads longer than 19,940 bp (Supplementary Data S1, Fig. S1A). We mapped^20^ all reads to human reference genome GRCh38^21^, and observed a median LRS aligned coverage of 17.2x (range: 10.0-94.3x, Methods, Fig. S1B) per individual. A median of 87.6% of basepairs aligned to the reference (Fig. S1C) and the median sequencing error rate was 11.6% (3.3% for insertions, 4.5% for deletions, and 3.8% for mismatches, Fig. S1D).

We generated a high-confidence SV set in four stages: (i) discovery, (ii) merging across individuals, (iii) genotyping, and (iv) imputation (Fig. 1A). We began (Fig. 1B) by discovering SVs with high sensitivity^8^ and refined them at predicted breakpoints using SRS data, when possible^22^ (Methods). Their presence was confirmed using the raw signal-level data^23^ (Methods, Fig. S6) to alleviate potential basecalling and alignment errors. We did not attempt to discover translocations and inversions. The SVs discovered across individuals were then merged and genotyped using two independent datasets: 3,622 and 10,000 Icelanders with LRS and SRS data, respectively^7,24^ (Methods). Finally, we imputed the genotyped variants into the long-range phased haplotypes of a total of 166,281 SNP chip-typed Icelanders^19,25,26^ and defined a set of high-confidence SVs, based on imputation accuracy and other filters (Methods).

**Fig. 1.**
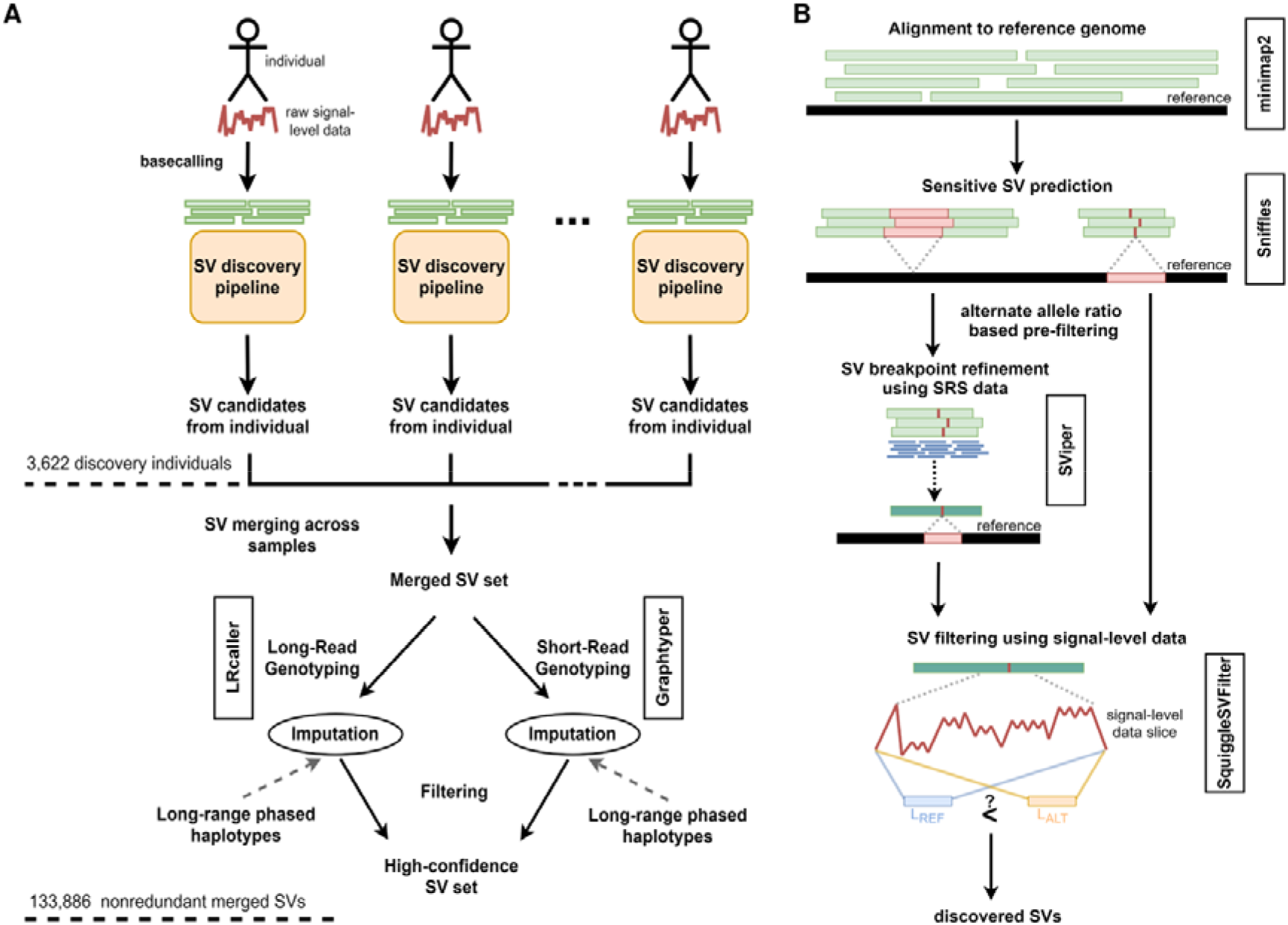
Structural variant (SV) analysis workflow. (A) Each individual is basecalled and their SVs are discovered independently. SV sets are merged across all individuals and the merged SV set is used to genotype individuals using both short read sequencing (SRS) and long read sequencing (LRS) data, separately. Finally, genotyped variants are imputed into long-range phased haplotypes and variants that pass SV filtering are accepted as high-confidence variants. (B) SV discovery pipeline: Reads are mapped to human reference genome (GRCh38) using minimap2, followed by the sensitive SV predictions using Sniffles. SV predictions are then pre-filtered based on their alternate allele ratio, and SV breakpoints are refined using SRS data, if possible, with SViper. Finally, candidate SVs are compared against the raw signal-level data using SquiggleSVFilter for further verification.

We identified 133,886 high confidence SV alleles (75,050 insertions, 55,649 deletions, and 3,187 unresolved insertions or deletions, Supplementary Data S2, Fig 2A), avoiding double counting of alleles of similar length at similar positions (Methods). We observed more insertions than deletions. This contrasts with results based on SRS^3,9^, where deletions are typically more frequent and easier to identify. We were able to impute 120,108 SV alleles (67,673 insertions and 49,845 deletions, 2,590 of unresolved insertions or deletions) into long-range phased haplotypes from 166,281 chip typed Icelanders. We identified a median of 22,636 SVs per individual (a median of 13,353 insertions and 9,474 deletions, Methods, Fig. 2A) of which a median of 20,891 were imputed, spanning a cumulative median length of 10.02 Mb per haploid genome. We estimated the false negative (FN) and false positive (FP) rates of our high-confidence SV set by comparing it to public SV datasets. Comparisons against an LRS SV dataset from Audano et al.^17^ (N = 15) and an SRS SV dataset gnomAD-SV^12^ (N = 14,891), using SVs within GiaB Tier 1 regions of HG002^5^, suggest FN rates of 2.6% and 3.4%, respectively, for our dataset (Methods). We estimated an FP rate of 8.2% for our dataset, by considering our common SVs absent in Audano et al and their observed versus expected allele counts in HG002. We also estimated FP rates of 6.3-7.6% for the gnomAD-SV dataset, which is comparable to the rate we estimated for our call set (Methods). These estimates may be upwardly biased due to population-specific drift. In an attempt to validate 70 of the SVs using PCR an SV was confirmed for 60 of the successful 63 assays (7 assays failed), suggesting a false positive rate of 4.8% (Methods, Supplementary Data S3).

**Fig. 2.**
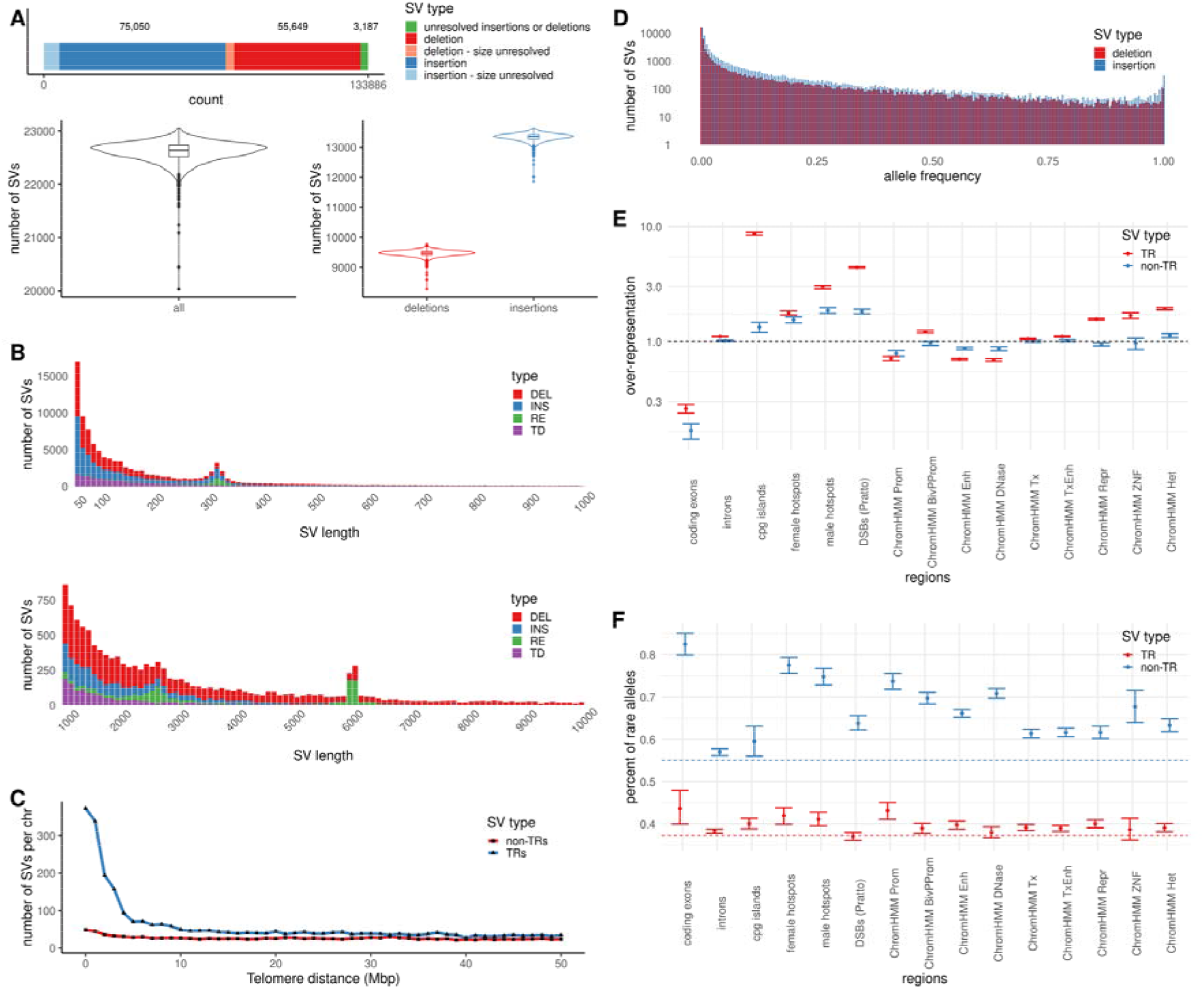
Merged structural variant (SV) set characteristics. (A) Number of merged SVs, and distribution of phased SVs per individual, total and stratified by SV type. (N = 3,204; 412 individuals not genotyped using short read sequencing (SRS) data, and 6 individuals in which < 90% of SV alleles could be phased, omitted. 6,455 insertions and 3,574 deletions are of unresolved size.) (B) Stacked SV length distributions in ranges [50 bp, 1 kb], and [1 kb, 10 kb], stratified by SV type. Insertions are subset into retrotransposable elements (RE), tandem duplications (TD). Remaining insertions are shown as INS. The peaks observed around 300 bp, 2.5 kb and 6 kb correspond to SINE, SVA, and LINE, respectively, highlighted by the increase in RE. (N = 120,670; 13,216 SVs without a specified size are omitted.) (C) Number of tandem repeat (TR) and non-TR SVs per chromosome as a function of telomere distance (binned at 1 Mb). (D) Allele frequency distribution of SVs binned at 0.05%. (E) Overrepresentation of SVs across genomic regions. (F) Percent of rare SVs across genomic regions. In panels E and F, values are mean and error bars indicate 95% confidence interval. (N = 120,670; 13,216 SVs without a specified size are omitted.)

To measure the relative merits of LRS versus SRS in SV discovery, we assessed whether SV alleles discovered using LRS were also found by gnomAD-SV^12^, which only uses SRS data. Comparing gnomAD-SV to the SV calls by Audano et al. with allele frequency greater than 50% suggested a 41.3% FN rate for the gnomAD-SV dataset (Methods). Similarly, among our set of 46,352 imputed SV alleles with frequency greater than 10%, 19,430 (41.9%) were not found in gnomAD-SV. Repeating this analysis for subsets of SV alleles within or outside of tandem repeat (TR) regions, we observed FN rates of 47.4% and 27.4%, respectively, for gnomAD-SV. LRS data also improved the genotyping of SVs in our data. Of 120,108 imputed SV alleles 76,857 (64.0%) and 3,917 (3.2%) SVs were imputed only from LRS and SRS genotyping, respectively. Furthermore, 74.2% and 38.6% of SV alleles within and outside of TR regions, respectively, could not be imputed using genotype calls from SRS. These results show that SV discovery and genotyping from LRS data is more accurate and reliable than from SRS^12^ data. The difference is particularly pronounced for SVs in TR regions, which have mutation rates 1 to 4 orders of magnitude higher than other genomic loci^27,28^.

The number of variants in our imputed SV set rapidly decreases with length (Fig. 2B), consistent with previous reports^12,15,17^. In order to better characterize insertions, we classified them into three groups; tandem duplications (TD), retrotransposable elements (RE), and other insertions (INS), corresponding to 30%, 7%, and 63% of insertions, respectively. We observe three noticeable peaks at sizes around 300 bp, 2.5 kb, and 6 kb, due to REs, corresponding to SINE, SVA, and LINE elements (Fig. 2B), as expected. We found more SVs, particularly TR SVs, near telomeres^17^ (Fig. 2C), a reflection of the sequence content of telomeres and the high mutation rate of TRs^17,29^. The number of alleles detected decreases with increased allele frequency, with 40.1% of them at a frequency less than 1%, i.e. rare (Fig. 2D, S2). The small number of variants that stand out for being fixed or near fixed in frequency are most likely examples where the reference sequence carries a derived rather than the ancestral state.

In general, frequency reflects the age of variants, such that younger variants are rarer than those that are older. As a result, differences in the relative allele frequencies of SVs by genomic region can provide information about the strength of negative selection that has acted against them. We observed both an underrepresentation of SVs and an elevated fraction of rare SVs in coding exons and non-coding regulatory regions, such as enhancers and promoters, compared to the genomic average (p < 0.002, in coding exons, enhancers, and promoters, for both TR and non-TR SVs, bootstrap test, Fig. 2E, 2F). We also found that SVs in TRs tend to be observed at higher frequencies than those outside of TRs (particularly when compared to those in coding exons), suggesting a higher tolerance of SVs in TRs. In regulatory elements, although SVs in TRs were more underrepresented than SVs outside TRs, they similarly had a higher fraction of common alleles than SVs outside TRs. In accordance with the notion that recombination plays a role in SV formation^30^, we found that SVs are enriched in double strand break regions^31^ and recombination hotspots^32^, particularly in TR alleles within male hotspots. (Fig 2E). Interestingly, we also observed an elevated rate of rare alleles in recombination hotspots (Fig 2F).

Variants inside coding exons that are not multiples of three in length generally result in translational frameshift and non-functional proteins. Among the 549 variants contained within a single coding exon, we observed a deficit in variant lengths that are not multiples of three, 187 (34.1%) compared to two thirds (362) expected (p = 4.9·10^−55^, binomial test, two-sided, Fig S3), in line with results using indels^26^. These results are consistent with the hypothesis that SVs that result in translational frameshift are selected against due to their phenotypic impact.

We used two strategies to determine the impact of SVs on phenotypes. First, we inferred association for SVs based on linkage disequilibrium (LD) with variants previously reported to be associated with phenotypes. Of the 116,479 unique variants reported in the GWAS catalog, 11,194 are in strong LD (R^2^ ≥ 0.8) with 5,238 SV alleles in our dataset, suggesting possible functional explanations for these associations (Supplementary Data S4). A subset of 34 high and 54 moderate impact SVs, overlapping exons or splice regions, are in strong LD with 198 GWAS catalog variants and are therefore plausible causal variants for the reported associations. Among these are examples where the presence of an SV was previously established using alternate methods, including a deletion in *LCE3B*^33^ associated with psoriasis, and a deletion in *CTRB2* associated with diabetes^34,35^ and age related macular degeneration^36^. Another example is a rare 2,460 bp deletion which removes two exons of *COL4A3* and associated with hematuria^37^. We also find loci where the occurrence of an SV at a GWAS locus has not been reported; including a deletion that overlaps the first exon of *SLC25A24* and is in strong LD with a SNP associated with white blood cell count^38^, and a 120 bp inframe deletion in *KAT2B*, that removes 40 amino acids from the translated protein and is in strong LD with a variant associated with systolic blood pressure^35,39^.

Second, we performed direct tests of association with phenotypes of studied Icelanders (Methods). We found an association with a rare 14,154 bp deletion overlapping the first exon of *PCSK9* (Fig. 3A) and LDL cholesterol levels (adjusted effect = −1.31 SD and p = 7.0·10^−20^, Fig. 3B). LDL cholesterol levels were 0.93 mmol/L lower in carriers (N = 75) than in non-carriers (N = 98,081). We observed 13, 56, and 119 heterozygous carriers of the deletion in our LRS, SRS, and imputation datasets, respectively, corresponding to an allele frequency of 0.037%. No homozygous carrier was identified. *PCSK9* encodes the enzyme proprotein convertase subtilisin/kexin type 9, a key regulator of LDL cholesterol metabolism^40^ and a target of cholesterol lowering drugs^41^. Loss-of-function variants in *PCSK9* are known to result in lower levels of LDL cholesterol and reduced cardiovascular risk^42–44^, consistent with the association observed here. We next tested this deletion for association with 4,792 plasma proteins measured in 38,405 Icelanders using SOMAscan^45^. PCSK9 levels of carriers (N = 20) were on average 1.99 SD below the population mean (p = 3.1·10^−13^, Fig. 3C). The geographical distribution of the carriers in Iceland suggests that the variant is more prevalent in Western Iceland than in other parts of the country (Fig. 3D). Two carriers were found in gnomAD-SV (1 out of 9,534 Africans and 1 out of 7,624 Europeans), and one carrier with low LDL levels was found in a study with Dutch participants^46^, showing that the deletion is not specific to the Icelandic population.

**Fig. 3.**
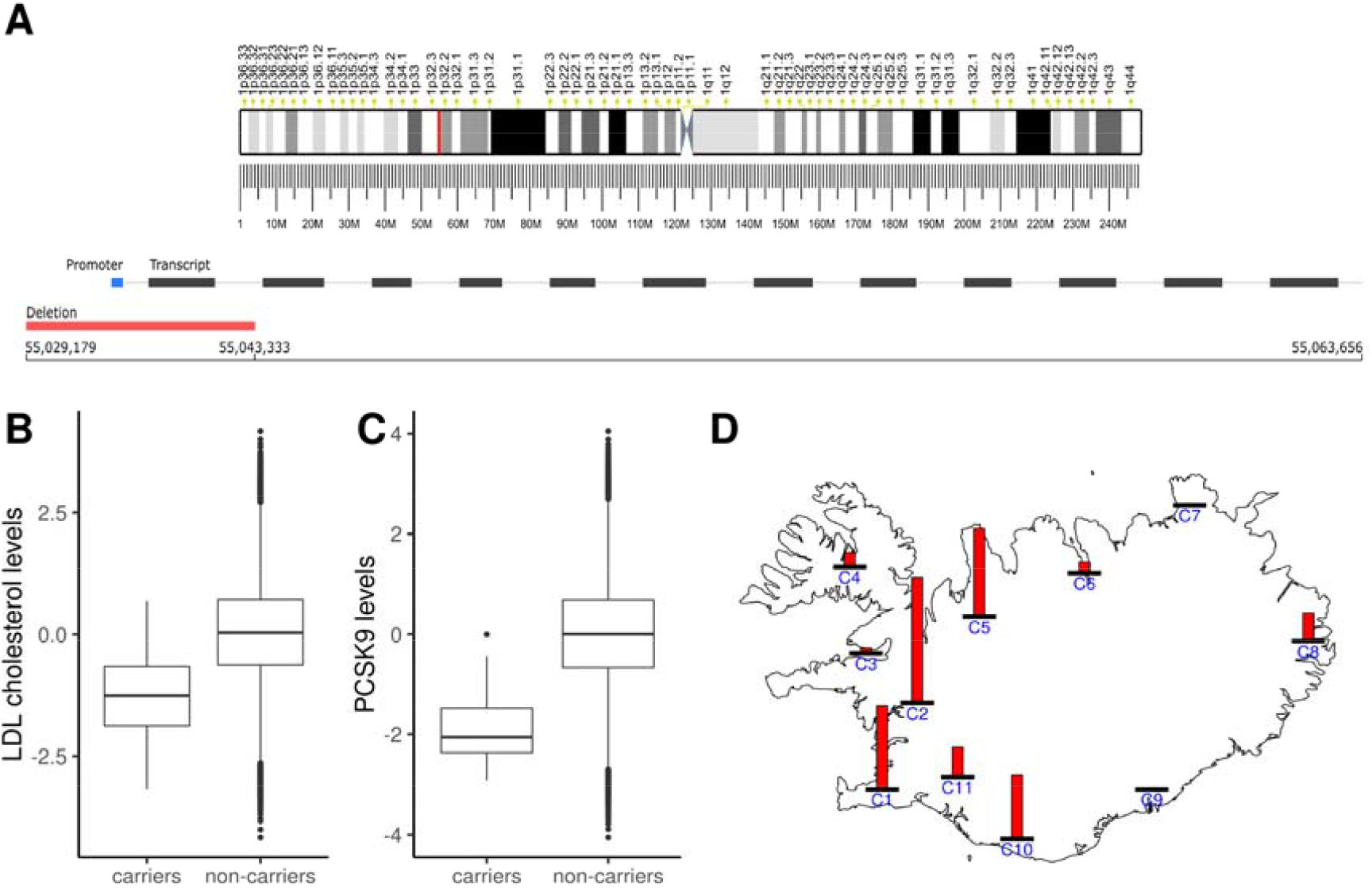
Large deletion in *PCSK9* associated with lower LDL cholesterol levels. (A) Chromosome 1 ideogram with cytobands, highlighting the SV site in red. A 14,154 bp (allele frequency 0.037%) deletion removes the promoter and the first coding exon of *PCSK9*, at basepairs 55,029,179 to 55,043,333 (GRCh38). (B) LDL cholesterol levels in carriers and non-carriers (n = 72 carriers, n = 96,840 non-carriers, effect = -1.31 SD, p = 7.0·10^−20^). (C) PCSK9 protein levels in carriers and non-carriers, using SOMAscan (n = 20 carriers, n = 38,385 non-carriers, effect = -1.99 SD, p = 3.1·10^−13^). (D) The geographical distribution of the *PCSK9* deletion in 166,281 chip-typed Icelanders. Each bar shows the allele frequency of the variant relative to the geographical region with the highest frequency. Icelandic counties are grouped into 11 regions C1-C11 as shown in the panel (see Methods, Table S5).

The SVs discussed above, although detected using LRS data, could also have been detected using SRS data, as they occurred in genomic locations where short reads can be reliably mapped. Below we present three examples of common multi-allelic SVs within tandem repeats, not found in gnomAD-SV dataset and are difficult to detect from SRS. These include the repeat region of an exon in *ACAN*, a proline-rich repeat region in *NACA*, and the zinc finger domain of *PRDM9*.

*ACAN* contains an exonic variable number tandem repeat (VNTR), with a 57 bp motif (19 amino acids in aggrecan, the translated protein), within one of its chondroitin sulfate (CS) attachment sites^47^ in which we identified 11 SV alleles; the reference allele along with deletions of 1, 3, 4, 5, 6, 8, and 14 motif(s) and insertions of 1, 2, 3, and 4 motif(s). (Fig. 4A, 4B). We found the 5-motif (285 bp) deletion to be highly correlated (LD R^2^ = 0.96) with a synonymous SNP (rs16942341[T], AF = 3%) reported to associate with decreased height (effect = -0.13 SD, p = 4·10^−27^) in a large GWAS analysis of 183 thousand individuals of European decent^48^. Both the reported synonymous variant and the SV are strongly associated with height in our data (effect = -0.13 SD, p = 1.02·10^−10^; and effect = -0.12 SD, p = 1.35·10^−9^, respectively). We observed a stronger association and a linear relationship (effect = 0.016 SD/motif inserted, p = 6.2·10^−18^, Methods) between the number of motifs carried and height (Fig. 4C, Table 1), suggesting that this SV plays a causal role in the association with height. The variance explained in height by the number of motifs carried (R^2^ = 9.9·10^−4^) is also higher than the variation explained by the SNP correlated with the 5-motif deletion (R^2^ = 5.7·10^−4^). The number of TRs results in a change in the number of CS attachment sites and thereby the number of attached CS chains on the aggrecan molecule^49,50^. Aggrecan is the most abundant proteoglycan in cartilage^51^ and the negatively charged CS chains have been shown to move water into cartilage^52^, thus the varying number of CS chains can affect the functional properties of this protein.

**Fig. 4.**
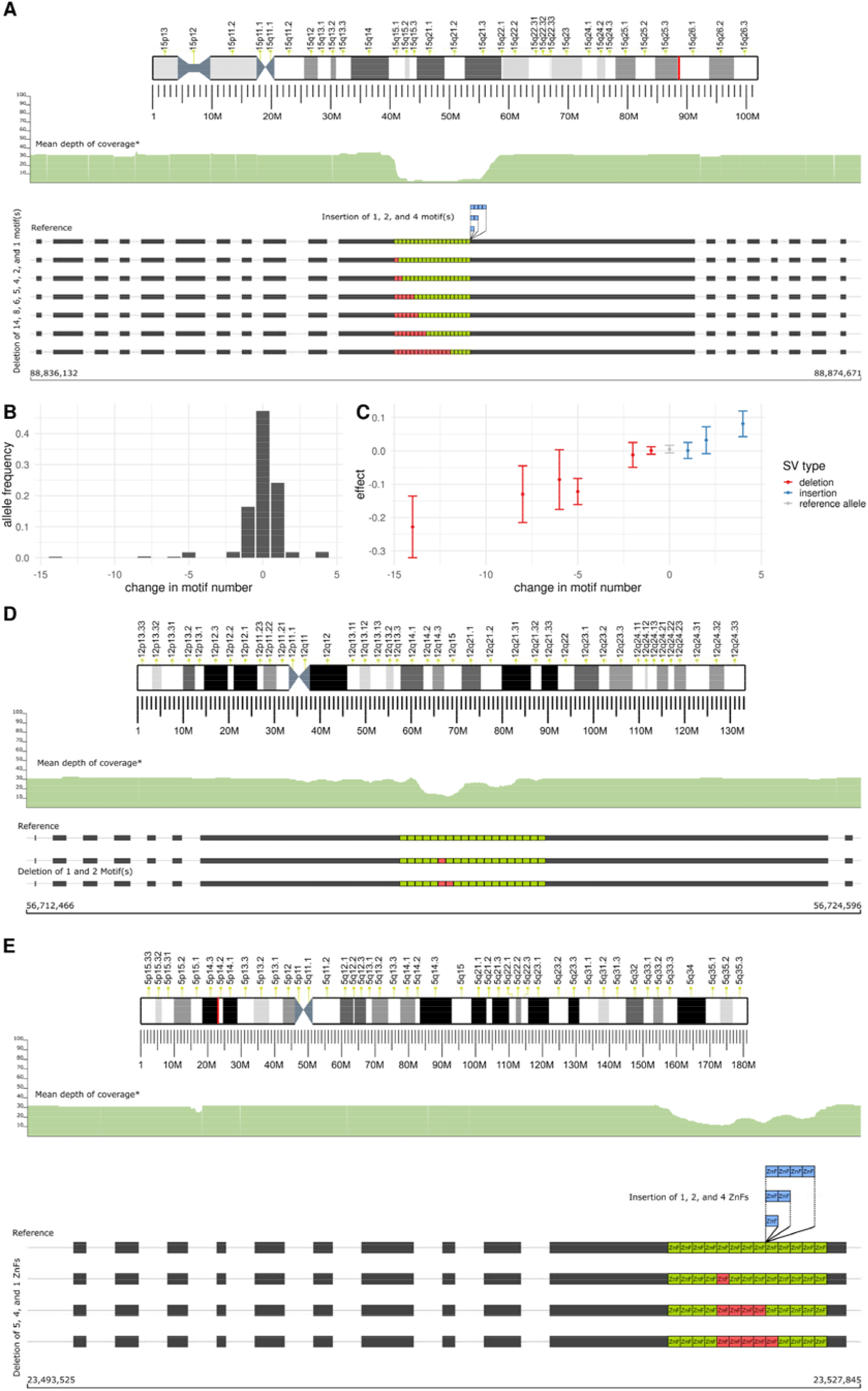
Multi-allelic SVs in repeat regions within exons of *ACAN, NACA*, and *PRDM9*, difficult for SV detection using short read sequencing (SRS). (A, D, E) Ideograms of respective chromosomes with cytobands highlight the SV site in red. *SRS mean depth of coverage tracks across respective genes are accessed via the gnomAD browser. Motif lengths, tandem repeat (TR) begin and end sites not to scale. Inserted or deleted motifs shown with arbitrary begin sites within the TR region. (A) SV alleles with inserted or deleted 57 bp motifs within an exon of *ACAN*. SRS coverage track shows the absence of reads with a reliable mapping across the TR region. (B) Allele frequencies of the SV alleles, including the reference allele. (C) Effects of the SV alleles and the reference allele on height, showing a linear relationship (effect = 0.016 SD/motif inserted, p = 6.2·10^−18^) Error bars indicate 95% confidence intervals (CI). SV alleles with frequency less than 0.01% omitted in panels A and B due to their large CI in effect. (D) SV alleles with deleted 69 bp motifs within an exon of *NACA*. (E) SV alleles with inserted or deleted 84 bp zinc finger motifs within the last exon of *PRDM9*. SRS coverage tracks in panels C, D indicate reduced reliably mapped reads across the TR regions, compared to the remaining of the gene.

**Table 1:** ACAN SV allele frequencies, lengths, p-values and effects on height, and SV types.

| freq | length | p-value | effect | event |
| --- | --- | --- | --- | --- |
| $3.0 \cdot 10^{-3}$ | -798 | $1.5 \cdot 10^{-6}$ | -0.228 | DEL |
| $3.9 \cdot 10^{-3}$ | -456 | $2.8 \cdot 10^{-3}$ | -0.130 | DEL |
| $3.2 \cdot 10^{-3}$ | -342 | $6.0 \cdot 10^{-2}$ | -0.086 | DEL |
| $1.7 \cdot 10^{-2}$ | -285 | $1.4 \cdot 10^{-9}$ | -0.122 | DEL |
| $9.8 \cdot 10^{-4}$ | -154 | 0.49 | -0.058 | DEL |
| $1.9 \cdot 10^{-2}$ | -114 | 0.53 | -0.012 | DEL |
| 0.17 | -57 | 0.86 | 0.001 | DEL |
| 0.47 | 0 | 0.40 | 0.005 | reference allele |
| 0.24 | 56 | 0.94 | 0.001 | INS |
| $1.8 \cdot 10^{-2}$ | 117 | 0.12 | 0.032 | INS |
| $7.4 \cdot 10^{-5}$ | 171 | $9.0 \cdot 10^{-2}$ | 0.504 | INS |
| $1.9 \cdot 10^{-2}$ | 228 | $3.6 \cdot 10^{-5}$ | 0.081 | INS |

*NACA* contains an exonic VNTR, with a repeat length of 69 bp (23 amino acids in NAC-alpha, the translated protein), repeated 18 times in GRCh38, in which we identified deletions of 1 and 2 motifs, (69 and 138 bp, respectively, Fig. 4D). The intergenic SNPs rs2860482[A] and rs7978685[T] have been reported to be associated with atrial fibrillation^53^ and both SNPs are in strong LD (R^2^ = 0.85) with the reference allele of the VNTR. This exon of *NACA* is transcribed in skNAC, a muscle specific, alternately spliced transcript whose importance in the developing heart has been demonstrated in animal models^54^. This suggests that the multi-allelic SV in NACA is the likely explanation of the observed associations with atrial fibrillation^54,55^.

Within the zinc finger (ZnF) domain of the recombination hotspot positioning gene, *PRDM9*, we identified deletions of 1, 4, and 5 ZnF(s), and insertions of 1, 2, and 4 ZnF(s), resulting in the removal or addition of a number of ZnF motifs to the encoded protein, PRDM9 (Fig. 4E). The ZnF domain of PRDM9 is the DNA binding domain and the SV alleles thus introduce alterations in the DNA binding motif of PRDM9 and consequently change the locations of meiotic recombination^56,57^. All the different zinc finger motif lengths show a strong association with the location of crossovers as measured by the fraction of crossovers that occur in recombination hotspots (Table 2). These results are consistent with previous results, which were indirectly ascertained via SNPs tagging multiple motif counts^32^, while the current results allow us to directly ascertain the effects of each motif count individually.

**Table 2:** PRDM9 SV allele frequencies, lengths, p-values (negative log) and effects on recombination hotspots, and SV types.

| freq | length | p-value (negative log) | effect | event |
| --- | --- | --- | --- | --- |
| $1.3 \cdot 10^{-2}$ | -420 | 11.541 | -0.204 | DEL |
| $5.6 \cdot 10^{-3}$ | -336 | 4.8091 | -0.201 | DEL |
| $2.3 \cdot 10^{-3}$ | -84 | 5.0098 | -0.318 | DEL |
| $9.1 \cdot 10^{-5}$ | 331 | 6.7388 | -1.679 | INS |
| $5.4 \cdot 10^{-3}$ | 168 | 304.31 | -1.700 | INS |
| $2.6 \cdot 10^{-2}$ | 83 | 1716.1 | -1.732 | INS |

We catalogued genes with rare homozygous loss of function (LOF) SVs, as predicted by Ensembl Variant Effect Predictor (VEP), and found 181 genes with a rare LOF SV, where at least one homozygous carrier was observed (151 of these were not found in the gnomAD-SV set, Supplementary Data S5). A number of these have been reported to cause diseases under recessive inheritance. One example is a 57 kb deletion, overlapping the genes *CTNS* and *SHPK*, originally associated with cystinosis^58^, a lysosomal storage disease characterized by the abnormal accumulation of the amino acid cystine, where homozygous carriers of the deletion generally develop cystinosis. We identified a single homozygous carrier of this deletion in our imputation set, who was not in our genotyping set and has been diagnosed with cystinosis. We also observed a 31.7 kb deletion (exons 11-17, AF = 0.45%) in *GALC* encoding galactosylceramidase. This is the most common mutation causing Krabbe disease in Europeans^59–61^. We identified a single homozygote for this deletion, a boy with a diagnosis of Krabbe disease.

In this study, we demonstrate the first application of LRS at population scale, and describe how it can be used to accurately identify SVs, and to assess their impact on human disease and other traits. We identified over 22,636 SVs per individual, three to five times more than found with SRS data^10,12^. We show that LRS is more sensitive than SRS in detecting SVs across the genome. This advantage is most pronounced in repeat regions, such as tandem repeats. We report 5,238 SVs in strong linkage disequilibrium with variants in the GWAS catalog that are associated with a disease or other traits. This constitutes an increase of over two-folds from the number of SVs using SRS data alone^12^. These results show that LRS can further our understanding of disease mechanisms and the effects of sequence variation on various human traits.

LRS technology and accompanying data analysis methods are still being developed and can be improved upon. Although we detected a large number of SVs per individual, we did not attempt to discover all forms of structural variation. We detected fewer long insertions than expected, possibly due to sequencing bias or limitations of the LRS analysis algorithms. We also expect an underrepresentation of very rare SV alleles as it is more difficult to phase and impute alleles with few carriers accurately. Although we have highlighted SVs that overlap coding exons due to their established functional impact, other SVs may still affect the individual, e.g. those altering regulatory regions or changing RNA secondary structure. A better understanding of the biochemical causes and consequences of SVs will be essential to understand human evolution and disease. These will in turn also lead to better analysis methods and increase our ability to identify SVs and assess their impact.

SVs have frequently been found using targeted approaches, often relying on discovered SNPs or indels in a disease association context. We demonstrate that our method can identify SVs in a genome-wide, non-targeted fashion. We show that SVs affecting protein function are disproportionately rare. As a result, large scale SV studies will be essential to characterize their role in the genetics of disease. This study, based on LRS data from 3,622 Icelanders, lays down an important foundation for further large-scale SV studies, allowing investigation of their full frequency spectrum, including those in genomic regions thus far inaccessible to SRS technologies.

## Acknowledgements

We thank Prof. Jared Simpson, our colleagues from deCODE genetics / Amgen Inc., and employees of Oxford Nanopore Technologies for their help and advice. We would also like to thank all research participants who provided a biological sample to deCODE genetics.

## Author contributions

DB implemented software with additional software implemented by HI, HPE, SK, SM, GH, and BVH. DB and BVH wrote the paper with input from HI, AO, HPE, EB, HJ, BAA, SK, MTH, SAG, RPK, GH, GP, OAS, AH, UT, HH, DFG, PS, OTM, KS. HI implemented the analysis pipelines, with input from DB, SK, SAG, STS, GM and BVH. DNM and OTM performed the ONT sequencing. AsJ, and AdJ performed PCR validation experiments. GE, IO and OS acquired LDL measurements. HH and BT acquired heart tissues. BVH and KS conceived and supervised the study. All authors approved the final version of the manuscript.

## Data availability

The SVs discovered in this study and correlations with GWAS catalog variants are included in the supplementary material. Icelandic law allows for unimpeded sharing of summary-level data. However, the law does not allow the sharing of individual-level data on genotypes and phenotypes outside of Iceland.

## Code availability

SViper, modified, used in this study: https://github.com/DecodeGenetics/SViper/tree/cornercasesScrappie, modified, used in this study: https://github.com/DecodeGenetics/scrappie/tree/v1.3.0.events

SquiggleSVFilter: https://github.com/DecodeGenetics/nanopolish/tree/squigglesv

LRcaller: https://github.com/DecodeGenetics/LRcaller

## Methods

### Participants

A set of 3,622 individuals was selected for ONT sequencing, including 441 parent-offspring trios. The individuals were selected from a large set of Icelandic samples collected as part of disease association efforts at deCODE genetics. The samples constitute a database of DNA sequence variation in the Icelandic population combined with extensive phenotypic data, including information on blood levels of lipids for up to 113,355 genotyped individuals (approval no. VSN-15-023). The study population has been described in detail previously^18,19,62,63^. All participants were Icelanders who donated biological samples for genotyping and provided informed consents as part of various genetic programs at deCODE genetics, Reykjavik, Iceland. The study was approved by The National Bioethics Committee of Iceland (approval no. VSN-15-023 and VSN 05-097, with amendments). A subset of individuals (N = 102) provided heart samples, with approval no. VSN-05-097.

### DNA source

Most of the DNA samples sequenced in this study were isolated from whole blood (N = 3,524). DNA from whole blood was extracted using the Chemagic method (Perkin Elmer), an automated procedure which involves the use of M-PVA magnetic beads (see URLs). DNA samples were also isolated from heart tissue (N = 102), 4 individuals provided both heart and whole blood samples. Samples were received and subsequently stored in liquid nitrogen. Samples were cut to smaller size on dry-ice if needed. Lysis buffer and a sterile 5mm steel bead were added to each sample prior to homogenisation on a TissueLyser LT (Qiagen). DNA was extracted from the homogenised lysates using the MasterPure DNA Purification kit (epicentre) following the manufacturers protocol but with overnight Proteinase K digestion. Isolated DNA samples were quantified using a Trinean DropSense™ and integrity assessed using the Fragment Analyzer capillary system from AATI.

### Sample preparation

Sequencing libraries were generated using the SQK-LSK109 ligation kit from ONT. Sample input varied from 1 – 5 µg of DNA, depending on the exact version of the prep kit and flowcell type used for PromethION sequencing. 1,322 of 4,757 flowcells underwent partial DNA shearing using the Covaris g-TUBE™ to a mean fragment size of 10 − 15 kb. The remainder of the samples were not sheared (Fig. S4 from March 2019 and onward). Library preparation started with DNA repair/A-tailing using the NEBNext FFPE repair mix (#M6630) and the NEBNext End repair/dA-tailing module (E7546), followed by AMPure XP bead clean-up. Adapter ligation was performed using the NEB T4 ligase (NEBNext quick Ligation Module, #E6056) and the ONT/LSK109 adapter mix (AMX) and ligation buffer, respectively. Samples were again purified using AMPure XP beads, using the Long Fragment Buffer (LFB) for the wash steps. Final sample elutions from the beads were done using 15 µL of elution buffer (EB). Samples were quantified using a Qubit fluorimeter and diluted appropriately for loading onto the flowcells.

### Sequencing

Samples were loaded onto PromethION R9.4.1 flowcells following ONT standard operating procedures. Sequencing was performed on PromethION devices. Data acquisition varied from 48 − 60 hours per flowcell.

### Basecalling

The squiggle data from the PromethION sequencers was basecalled using Guppy (3,622 individuals, 4,757 flowcells). We ran Guppy Sequencing Pipeline Software for GPU machines, version 3.2.2 (643 flowcells) and version 3.3.0 (4,114 flowcells), using the *flipflop* model (configuration file `template_r9.4.1._450bps_large_flipflop.jsn` from guppy version 2.3.1) for PromethION flowcells with firmware 2.0.4 and the *hac* model (using the corresponding configuration file `template_r9.4.1._450bps_hac_prom.jsn`) for firmwares 2.0.10, 2.0.12 and 2.0.14 (Fig. S4).

Our oldest flowcells were originally basecalled using Albacore (a now deprecated basecaller from ONT). Mid-year 2019 we upgraded to Guppy version 3.2.2. This was reported in our preliminary study based on 1,817 individuals^64^. In autumn 2019, we upgraded Guppy to version 3.3.0 due to new PromethION firmware, and re-basecalled all flowcells previously basecalled with Albacore to reduce error rates^65^.

All 3,622 individuals basecalled with Guppy had a minimum reference genome aligned sequencing coverage of at least 10x at the time of analysis for SV discovery.

### Read mapping

The basecalled reads were mapped to human reference genome GRCh38^21^ with minimap2^20^ (version 2.14-r883), using the recommended option for ONT sequence to reference mapping `(-x map-ont`). In addition we used the parameters `--MD -Y`. The aligned reads were sorted using samtools sort^66^ and stored in a BAM file.

### Sequencing statistics

To estimate sequencing error we used the following terminology:

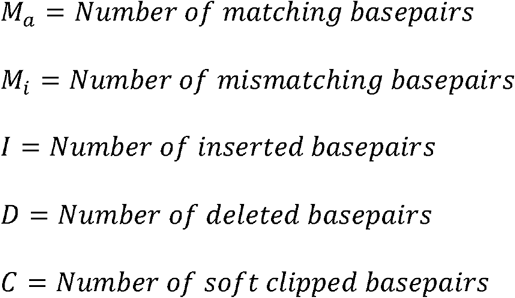

Then we have the quantities

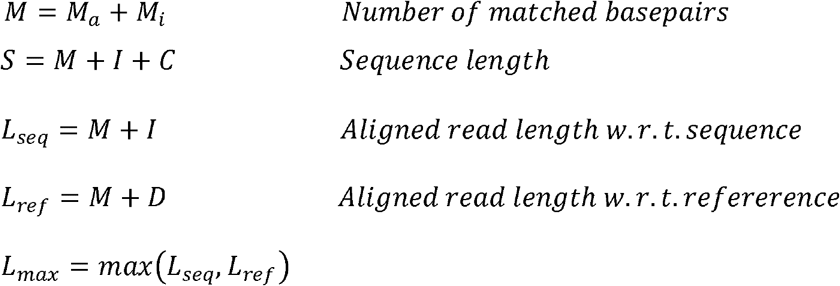

All secondary alignments are ignored. Quantities *M*_*i*_, *I, D, L*_*seq*_, *L*_*ref*_ and *L*_*max*_ are summed over all non-overlaping supplementary alignments for a read, if they exist. These quantities are calculated for all reads longer than 3000 bp for each basecalled flowcell. Shorter reads often have an ambiguous mapping to the reference genome and are therefore of limited use for SV calling. We did not omit reads labelled as “FAIL”, omitting those reads from analysis would result in lower error rates, higher mapping rates but lower sequencing coverage. Let *r* be a read and *R* be a set of reads. We report the error rate *E* as the total sum of erroneous basepairs, normalized by alignment length,

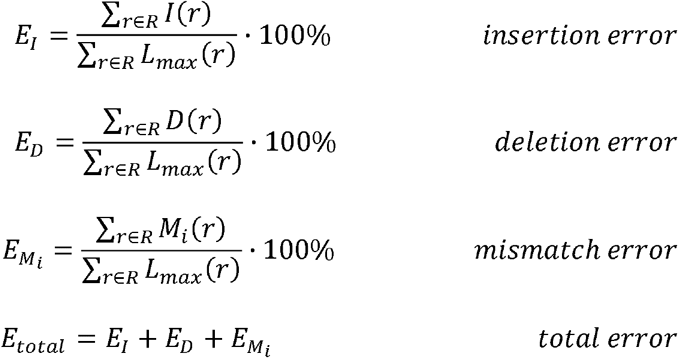

and the alignment accuracy, A, as alignment length normalized by its read length,

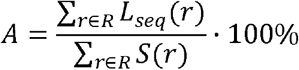

In addition we report aligned coverage as:

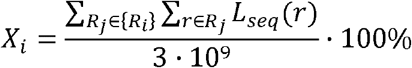

For all reads in a set of flowcells R_i_ belonging to the same individual i.

### Generating SV candidates from an individual

As shown in Fig 1B, after basecalling the raw reads, we align them to the GRCh38 reference genome using minimap2, and perform a sensitive SV prediction using Sniffles. We filter the SV candidates using their alternate allele ratios, and refine their breakpoints using SViper^22^. Next, we use SquiggleSVFilter on both the breakpoint refined SVs and filtered Sniffles SV calls (breakpoint unrefined) to verify their presence using the raw signal-level data. Fig. S5 shows the number of discovered SVs per individual, sorted by the aligned coverage of the individual.

### Preliminary structural variant predicting with Sniffles

A set of preliminary variant predictions is obtained using Sniffles^8^ (version 1.0.10) for each genome, in a highly sensitive fashion (using `-s 3`, and `--ignore_sd`) to minimize false negatives due to the existence of low coverage regions. Up to 30 supporting reads are reported per variant. Other optional parameters are left as default. Insertions and deletions with different start/end chromosomes and larger than 1 Mb are discarded. Next, deletions and insertions with alternate allele ratio below 0.2 and 0.05, respectively, are discarded as a pre-filter from raw Sniffles calls. A higher value was used for deletions as the basecaller is deletion biased. We calculate alternate allele ratio as the number of reads supporting the variant divided by the coverage at the variant site.

### Breakpoint and variant refinement with SViper

The variant predictions are breakpoint refined with SViper^22^ (see URLs, Code availability) using SRS data, when possible. All optional parameters were left as default. SViper first identifies the ONT reads supporting a candidate variant and forms consensus sequences flanking the candidate breakpoints. It then selects SRS reads near the predicted SV breakpoint(s), if available, and aligns to the ONT consensus sequence for polishing. The polished consensus is re-aligned to the human reference to provide the refined breakpoints.

### SV filtering using signal-level raw Nanopore data (SquiggleSVFilter)

We developed SquiggleSVFilter (URLs) to filter false SV predictions using the signal-level raw ONT sequencing data, i.e. the squiggle. SquiggleSVFilter employs the squiggle-vs-sequence log likelihood score function provided by Nanopolish^23^, and compares the log likelihood scores of the predicted alternate allele vs. the reference allele on the squiggle around both of the SV breakpoints. The likelihood score is essentially the probability of the signal-level raw data given a candidate sequence^23^. Nanopolish uses the events, which are the step-wise changes in the measured electrical currents, as the signal data in its log likelihood score function. Accessing an event interval over a predicted SV breakpoint requires a mapping of the read sequence indices to reference genome coordinates (i.e. a reference aligned BAM file) and to event indices, called an *event table*. To achieve this, we generated basecalls and event tables for reads that support a predicted SV, using a modified version of Scrappie (URLs, Code availability), and mapped these reads to the reference genome, using minimap2 with parameters as described in “Read mapping”.

For any SV supporting read, reported by Sniffles, we start by calculating the locations of the SV breakpoints in the read using the BAM file (Fig. S6). Next, we determine the read regions spanning the SV breakpoints, which we refer to as *subreads*. An alignment may not contain an anchor on the reference on both sides of a breakpoint, and may instead be soft-clipped on one of the sides. We therefore determine a left and right subread by approaching from the left and right flanks of the variant. Fig. S6A and S6B, depict sample deletions and insertions, respectively.

We define *B*_1_*B*_2_ as the left and right breakpoints from the reference (i.e. alignment anchors) for a SV. Note *B*_1_ = *B*_2_ for insertions. Using a flank size of 500 bp in the reference, we compute the read indices of the left and right subreads as follows,

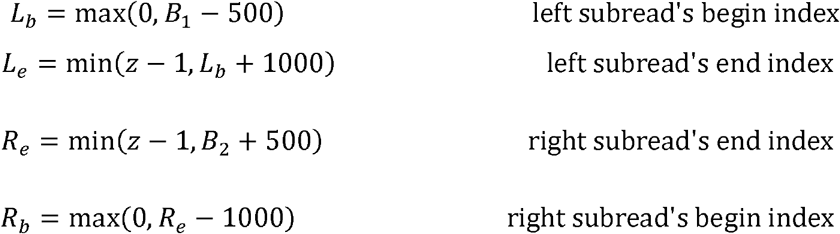

where is the length of the read.

Using the event table, we find the event indices corresponding to the subreads to access event slices spanning the SV breakpoints. Given the predicted SV breakpoint sites and sequence (for insertions), and the BAM file, we determine the reference (ref) and alternate (alt) allele sequences spanning the left and right subreads. We set the ref allele sequences as 500 bp of reference sequence flanking (*B*_1_ = *B*_2_) from both sides, for the left and right event slices, respectively, namely:

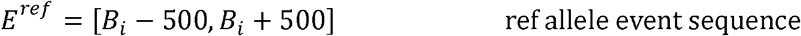

For deletions, *i* = 1 for the left breakpoint and *i* = 2 for the right breakpoint (shown as the blue/yellow and purple/red sequences in Fig. 6A). For insertions, *B*_1_ = *B*_2_ therefore 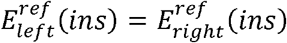 (shown as the blue/yellow sequence in Fig. 6B). We set the alt allele sequence as 500 bp flanking *B*_1_ from the left, followed by 500 bp flanking *B*_2_ from the right, for both deletion event slices, namely,

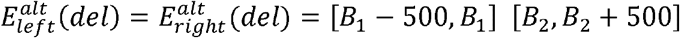

where denotes concatenation (shown as the blue/red sequence in Fig. 6A). For insertions, we set the alt allele sequence as 500 bp flanking *B*_1_ = *B*_2_ from the left or right, appended by the insertion sequence of the predicted SV. Namely, the alt allele event sequences are

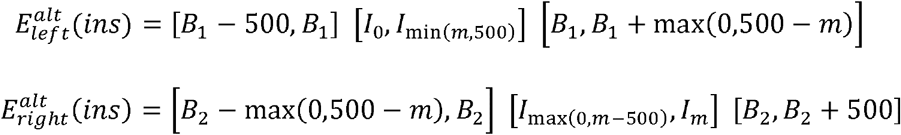

where is [*I*_0_, *I*_*m*_] the insertion sequence with length *m* (shown as the blue/purple/red, and purple/red/yellow sequences in Fig 6B, where m is depicted to be greater than *m*).

Finally, we calculate the raw signal-vs-sequence log likelihood scores using the ref and alt allele sequences for both event slices, and use their difference to support or reject the candidate variant. We support a variant if at least 3 reads obtain a log likelihood score difference of at least 1.92 for either of the event slices.

### Precision and recall in SV discovery

In order to estimate precision and sensitivity of SV discovery, we ran our pipeline with a subsampled version of the annotated HG002 dataset^67^ (aligned coverage of 16.3x), to reflect the coverage trends we obtained in our study. We define an SV in dataset A is “found” in dataset B, if there exists an SV in dataset B within 500 bp to the SV in dataset A, using their begin sites. We estimate a precision of 95.3% using discovered SVs within the Tier1 benchmark region, where 9,272 of 9,728 SVs with a unique begin site can also be found in the HG002 dataset. This rate is 82.9% (18,244 of 22,004) using all discovered SVs. We observed lower precision prior to using SquiggleSVFilter where 93.2% of Tier1 SVs (9,371 of 10,057) and 79.1% of all discovered SVs (18,835 of 23,819) can be found in the HG002 dataset. We estimate a sensitivity of 70.7 and 73.9% (9,558 and 9,992 out of 13,524) within the Tier1 benchmark region, using our discovered SVs larger than 50 bp, and using all discovered SVs (larger than 30 bp, default length value by Sniffles), respectively. We observed comparable sensitivity values prior to using SquiggleSVFilter, where 71.1% and 74.4% (9,618 and 10,061 out of 13,524) of the HG002 Tier1 SVs were found.

We observed SquiggleSVFilter to be especially effective in individuals with high number of SVs after the alternate allele ratio based pre-filtering, a result of high error rates in some of the LRS data. Fig. S7 shows this uneven level of filtering achieved by SquiggleSVFilter, across individuals, which is a result of the high mean and variance observed in error rates (Fig. S1D). Out of a total of 85M Sniffles-prefiltered SV calls, we retained 76.4M (89.8%) after using SquiggleSVFilter. In our previous study^64^as discussed in “Basecalling”, out of a total of 22.6M Sniffles-prefiltered SV calls, we retained 12.7M (56.4%) after using SquiggleSVFilter. The decrease in the percentage of filtered SVs is expected as in our current dataset we re-basecalled our raw reads using Guppy, a basecaller with lower error rates, resulting in less false positive SV calls, compared to Albacore.

We computed the percent of child SVs not found in any parent in 441 parent-offspring trios (Fig. S8). This percentage captures the rate of false positives among all positives in the child, as well as false negatives in the parents. On average, the percentage reduces from 4.5% to 2.6% after using SquiggleSVFilter, including samples with up to nearly 20% of reduction (from 22.7% to 2.9%) in this percentage. After applying SquiggleSVFilter we observed more comparable percentage values across samples, which we expect to limit the accumulation of false positive SV calls when merging variants across large sample sizes.

### Filtering individuals

In order to filter samples that may have a high number of false positive SVs, we compared the accepted SV sets from individuals to the SV set by Audano et al.^30^ We filtered an individual if less than 80% of their SVs (either breakpoint refined or unrefined) are found in the set by Audano et al. As a result, 43 individuals are not included in our SV analysis. Both breakpoint refined and unrefined SV sets from the remaining 3,622 individuals are moved to the SV merging step.

### SV merging

Most SVs are carried by multiple individuals, and thus will be re-discovered, potentially with slightly different representations across carriers, varying in length and location. In order to eliminate such redundancies, we applied the following SV merging approach, in which we represent SVs as vertices in a graph and find cliques representing merged SVs.

i. **Determination of tandem repeat (TR) SVs**: SVs occuring within a TR region may be assigned arbitrary breakpoints by the mapper and the SV discovery method, due to their repeat structure, and the error rates observed in ONT sequencing reads, which may prevent mappers from exploiting the small differences in the repeats, if they exist. We therefore first separate SVs into TR and non-TR and use different merging strategies for each set. We ran Tandem Repeats Finder^68^ on the reference genome (parameters 2, 7, 7, 80, 10, 22, 1000 for Match, Mismatch, Delta, PM, PI, Minscore, and MaxPeriod, respectively); in order to extract the TR regions from GRCh38. Next, we intersect the SV breakpoints with the TR regions and find the SVs contained within TR regions. Since an SV can reside within multiple TR regions, we assign the largest one to the SV.
ii. **Pre-clustering SVs prior to merging**: We pool all SV candidates, separately per chromosome and event type (insertions and deletions), and cluster them into disjoint SV groups prior to finding SV cliques to reduce the input sizes. We move the begin site of all TR SVs, to the begin site of their respective TR region. This pre-clustering iteratively finds gaps first in SV positions, then in SV lengths, such that no clique can be formed between two separate SV groups. We start by initializing our SV groups as a single SV group containing all SVs. For each SV group, we first sort SVs by their begin site, and split them into smaller non-overlapping groups, such that no SV in group A overlaps with any SV in group B. Next, for each SV group, we sort SVs by their length, and similarly split them into smaller groups, where two adjacent SV have a *distance* more than *D*, regardless of their begin site. We repeat these two steps until no SV group can be split any further. We define distance as,

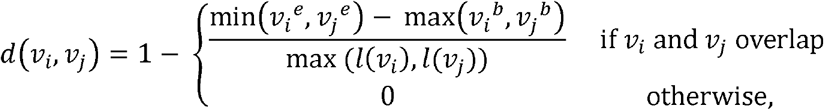

Where *υ*_*i*_ and *υ*_*j*_ represent two SVs, and *l*(*υ* _*i*_) the length of *υ* _*i*_. *υ* _*i*_^*e*^ and *υ*_*i*_ ^*b*^ represent the end and begin site of *υ*_*i*_. We measure the overlap between two insertions by representing them similar to deletions, where an insertion end site is set as its begin site incremented by its length. We chose *D* as 0.5.
iii. **Finding SV cliques**: We find cliques within each detached SV group generated in **ii**, independently for TR SVs and non-TR SVs as defined in i. We use an undirected graph *G*(*V, E*) where each vertex *υ ∈ V* represents an SV, and each edge *e ∈ E* between vertex *υ* _*i*_ and vertex is *υ*_*j*_ drawn if is at most *D*, and assign *d*(*υ* _*i*_ *υ j*) as the edge length. We then formulate the SV merging as a corrupted cliques problem, where given a graph, the aim is to transform into a clique graph with the smallest number of edge additions and removals, such that a clique represents a single merged SV. To solve this, we employ the Cluster Affinity Search Technique (CAST) algorithm^69^. We compute the average distance of vertex *υ* _*i*_ to the cluster as a weighted average such that

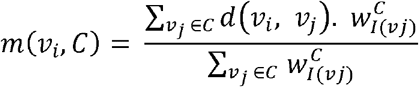

where I(*υ* _*j*_) is the *υ* _*j*_ individual the is discovered in, and 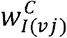 is the weight of *υ* _*j*_ for cluster *C*, set as the inverse of the number of SVs from individual *I*(*υ* _*j*_) found in the cluster *C*. The weighted average limits bias in the clique formation towards individuals with multiple SV calls at similar positions (detected with different approaches, e.g. breakpoint refined and unrefined SV calls). Likewise, the degree of a vertex *υ*_*i*_ is also calculated as number of individuals *I*(*υ* _*j*_) it has an edge with. We use *D* = 0.5 for non-TR SVs as described in **ii**, and *D* = 0.15 for TR SVs. We decided to be more conservative in the merging of TR SVs compared to non-TR SVs in order to be able to distinguish alleles with different number of repeats within the TR regions, and because we artificially increased their overlap by changing their begin sites as described in **ii**.
iv. **Finding SV clique representatives**: We represent a clique using an SV with the most common (begin site, length) attribute set among all the clique SVs. If there is no such single most common attribute set, we sort the SVs with the most common attribute sets using the frequency of their begin/end site, and length, among all the clique SVs, separately, in the given order. In this step, we use the original begin/end sites for TR SVs. For all TR SV clique representatives within the same TR region, we assign their position as the most common original begin/end site among all SVs of all cliques within the TR region. All further ties are first broken by prioritising begin sites over end sites, then using the alternate allele ratios. If ties cannot be broken, the median begin site is used.

SV clique representatives are finally presented as merged SVs. We note that variants discovered with length between 30bp and 50 bp are not filtered out during the SV merging step in order to prevent clique formation with incomplete data.

### Creating a Variant Call Format (VCF) file for the merged SV set

We constructed a VCF file using the SV clique representatives. We combined the SVs starting or ending at the same site into multi-allelic variants, and added the remaining variants as bi-allelic variants. We refer to this set of variants as *primary form* variants. We also added all multi-allelic variant alleles as separate bi-allelic variants, and we combined bi-allelic or multi-allelic variants that reside within 250 bp of each other as additional multi-allelic variant alleles; both combined, and separately for insertions and deletions. We refer to this second set of variants as *secondary form* variants. As when merging SVs, we did not filter out variant alleles of multi-allelic variants less than 50 bp. We also record a *TR region* for TR SVs and a *variant region* for multi-allelic SVs. A TR region contains the genomic coordinates of the begin and end sites of the TR an SV is found within. A variant region covers the minimum and maximum genomic coordinates of all the SV alleles used in a multi-allelic SV.

### Individual selection for short read genotyping

We selected 10,000 individuals for short read genotyping from our set of Illumina WGS individuals generated previously at deCODE genetics^19^ from the most recent freeze available. This set included 3,210 of the 3,622 ONT sequenced individuals. To increase the probability of finding multiple carriers of very rare variants, we added the parents of the ONT sequenced individuals which did not already exist among the 3,622 individual set, amounting to an extra 1,262 individuals. The remaining SRS individuals were selected at random from a set of 49,962 sequenced individuals.

### SRS genotyping with GraphTyper

We provided the merged SV set to GraphTyper^7,24^ v2.6, which generates an augmented graph genome using the SV predictions, together with previously discovered SNPs and indels^7^, for population scale genotyping. The variants were genotyped on the set of 10,000 SRS individuals, using three genotyping models for deletions and insertions. For insertions, we use the models: first breakpoint (B1), second breakpoint (B2), and their aggregate (AG). For insertions, we use the models: breakpoint (B), coverage (C), and their aggregate (AG). Since GraphTyper does not support multi-allelic SV genotyping, we added the multi-allelic SVs as separate bi-allelic variants.

### LRS genotyping (LRcaller)

LRcaller is a proof-of-concept genotyping algorithm that genotypes SVs directly from ONT sequencing reads. We introduced LRcaller v0.1 in our previous study^64^ where we genotyped 1,817 LRS individuals. In this study, we introduce LRcaller v0.2 which now allows for the genotyping of multi-allelic variants, better treatment of tandem repeats and additional genotyping models compared to v0.1.

Each breakpoint is genotyped independently, resulting in two sets of genotypings for the canonical deletion and insertion variants identified in this study, corresponding to the left and right breakpoints (Fig. S6). Note that the algorithm processes each variant independently, i.e. each variant is genotyped without considering other variants in the region, which may lead to suboptimal behavior when there are multiple neighboring variants.

In order to capture multiple types of information that could represent an SV, we use five genotyping models: direct (AD), variant alignment (VA), reference aware variant alignment (VAr), presence (PR) and joint (J). We use the reads overlapping a breakpoint and two sets of evidence for genotyping; (AD) from an alignment of a subread to the reference and alternate alleles and (VA, VAr, PR) from the alignment present in the BAM file as aligned by minimap2. The joint model (*J*)uses both sets of evidence.

#### AD genotyping

We start by constructing a sequence for the reference and the alternate allele from a VCF record, in a method analogous to the one described by SquiggleSVFilter. A sequence for the reference allele is constructed as the sequence in reference coordinates (*b* − 500, *b* − 500), where *b* is the reference position of the breakpoint. A sequence for the alternate allele is constructed analogously, except the sequence in the ALT field of the VCF record is first inserted into the reference and the sequence of the REF field is removed, creating a new reference ℛ. The alternate sequence constructed is the sequence in coordinates (*b* − 500, *b* − 500) in ℛ.

Next, we select and crop reads used for genotyping. For left breakpoints we select reads that map to a position in the interval, (*b* – 500, *b*). We let *c* ^L^ be the smallest position in the interval, (*b* – 500, *b*), where the read is aligned, let *i*(*c* ^*L*^) be the read index aligned to reference position *c*^*L*^ and we crop subsequence

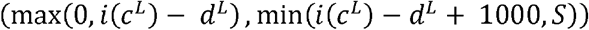

from the read, where *d* ^*L*^ = *c* ^*L*^ − *b* + 500, and *S* is the read length. For right breakpoints we select reads that map to a position in the interval (*b,b*. + 500) We let *c*^*R*^ be the largest position in the interval (*b,b*. + 500) where the read is aligned and we crop subsequence

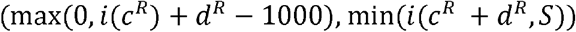

from the read, where *d*^*R*^ = *b* + 500 − *c*^*R*^. Cropped sequences shorter than 500 bp are ignored.

We then align cropped reads using the function globalAlignment in Seqan^70^, with scores 1 for match and -1 for mismatch, insertions and deletions, to the reference and the alternate alleles, producing alignment scores *q*(*a* _*i*_), for each allele *a* _*i*_ Reads where *q*(*a* _*i*_) is constant for all alleles are not used, and alignment scores less than 400 are artificially set to 400 in order to only use reads that map to at least approximately half of either the reference or the alternate allele. For all other reads, we first identify an allele *a*_max_ with the highest alignment score and then compute a score

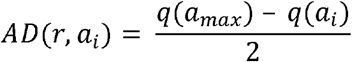

for each allele *a*_*i*_. We cap *AD* (*r, a*_*i*_) to have value at most 10.

#### VA, VAr, PR genotyping

We consider the minimap2 alignment to the reference in an interval near the SV breakpoints. This window is determined as

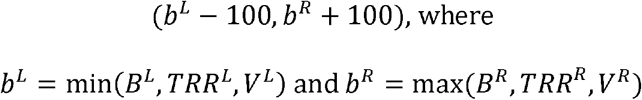

such that, (*B* ^*L*^ and *B* ^*R*^),(*TRR* ^*L*^,*TRR* ^*R*^) and (*V* ^*L*^, *V* ^*R*^)represent the SV breakpoints, tandem repeat region and variant region, respectively. Tandem repeat regions are only identified in variants within tandem repeats and variant regions are only defined for multi-allelic secondary variants. When considering the left breakpoint we only consider reads overlapping reference coordinate *b* ^L^− 100 and are not soft clipped (> 500 bp) at the start of the alignment. An analogous exclusion criteria is applied when the right breakpoint is considered. considered to support the alternate variant. For other reads we calculate L, the number of Reads that are soft-clipped (> 500 bp) at the end of their alignment within this window are inserted minus the number of deleted basepairs, occurring in stretches of at least 5 bp, in this window.

We then use three different models to turn these alignment information into genotypes; VA, VAr and PR. In the *VA* model we start by sorting all alleles by their length and compute their distance from the reference. We then determine the allele *a* _e_ with length distance from the reference closest to *L* and assign

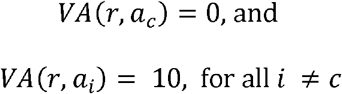

*VAr* is a reference aware model and is only used for bi-allelic variants, reads where 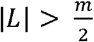 support the alternate allele and those where 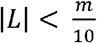 support the reference allele, where *m* is the SV length. Reads supporting the alternate allele are given a score

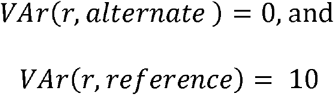

while those supporting the reference allele are given scores

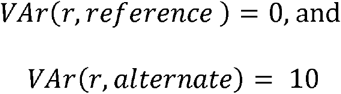

Reads supporting neither the reference nor the alternate allele are not used. The PR model only checks the presence of an SV within the region, regardless of the variant called. A bi-allelic call is made, for a reference and alternate allele, reads are assumed to support the reference or an alternate allele when |L| < 30 and |L| ≥ 30, respectively. Reads supporting an alternate allele are given a score

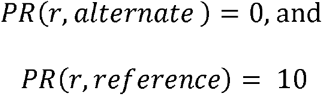

while those supporting the reference allele are given scores

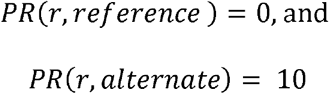

#### Joint model genotyping

The output of the AD and the VA models are then used in a joint model as:

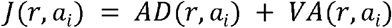

#### Relative log likelihoods

These scores are then used to compute *I* (*r*| *a*_1_ *a*_2_), the relative log likelihood of observing a read given a genotype, represented by a pair of alleles (*a*_1_ *a*_2_.) A score *f* (*a*_1_,*r*) (where *f* is one of AD, VA, VAr, PR, J) is interpreted to represent relative likelihood in log2 scale of observing the read *r* from a haplotype containing the allele *a*_i_. Assuming *f* (*a*_1_,*r*) ≥ *f* (*a*_2_,*r*), We let

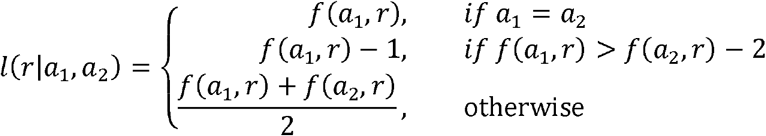

A joint relative likelihood of observing all the reads given the three possible genotypes is then found by multiplying these relative likelihoods, or summing the log relative likelihoods.

### Genotyping with LRcaller

The variants were genotyped independently for the left and right breakpoint using the five different models presented above for each variant, producing a total of ten genotypes per individual/marker pair.

### Evaluation of LRcaller

i. **Simulations:** To evaluate the ability of LRcaller to call SVs we simulated SVs by creating a new reference genome and mapping the reads of an individual to this new reference. We started with an earlier set^64^ of 68,050 SVs. We randomly selected 9 samples and for each sample we performed 10 simulations. We looped over all SVs in the VCF file of 68,050 SVs and created a corresponding SV in a new reference genome, with the roles of insertions and deletions reversed. We split the SVs into two categories, tandem repeat (TR) and non-tandem repeat (non-TR). For TR SVs we randomly selected a location for the SV among all tandem repeats of length equal or greater to the SV being studied, we then inserted or deleted a sequence at the location of the tandem repeat with length equal to the length of the SV. Inserted sequences consisted of the motif of the TR at the location of insertion/deletion. For non-TR SVs we selected a position randomly from the reference genome to be the start site of a deletion, taking care that this position was not annotated as ‘N’ and it was not within 100 bp from a tandem repeat. At this location we inserted the sequence of the deletion or deleted a sequence of the same length as the inserted sequence. We then mapped all reads to this new reference and genotyped variants. We finally measured SV calling accuracy as our ability to recall the variants at these sites. We recalled 92.3% of the variants using at least one of the genotyping methods, implying an estimated 7.7% false negative rate for LRcaller. A subset of these false negatives may be due to true SVs in the sample at the simulated location, mapping artefacts and low coverage at the variant site.
ii. **Additionally ONT sequenced individuals:** Following the generation of the LRS dataset of 3,622 individuals analyzed in this study, we sequenced an additional 491 individuals. Using 117,108 high-confidence SV alleles genotyped with an LRcaller model (excluding those genotyped with Graphtyper using SRS), we compared the genotypes of these individuals as output by LRcaller to the genotypes from imputation. We observed a mean and median error rate of 2.8% and 0.42%, respectively. When considering high quality rare variants our mean and median error rates were 0.66% and 0.0%, respectively. As expected, the error correlates with imputation info and phred scaled genotype likelihood. Restricting our analysis to alleles with imputation info above 0.99 and allele proband pairs where the phred-scaled genotype likelihood was above 40 give a mean and median error rate of 0.45% and 0%, respectively. Genotyping error rate per marker in the extra sequenced samples is given in Supplementary Data S2.
iii. **Comparison to HG002:** We genotyped the individual HG002 sequenced to 108.1X and basecalled using Guppy 3.3.2. As this coverage was higher than the one used in our study and to see the effect of coverage on we also subsampled the file to 0.1, 0.2, 0.4 and 0.8 times its original coverage. Subsampling was performed using samtools view -s a.x, where x is 3.1,2.2,1.4, or 0.8, for coverage of 0.1,0.2, 0.4 and 0.8, respectively. We first used LRcaller to genotype variants represented by the GiaB for HG002 (HG002_SVs_Tier1_v0.6.vcf)^67^. These variants were ascertained in GRCh37 and were liftover to GRCh38 using gorpipe^71^, variants where both the reference and alternate allele were shorter than 50bp were removed, leaving 30,278 variants for analysis.

Genotyping using LRcaller resulted in HG002 being called as a carrier of 28,935 (95.7%) of the variants, suggesting a false negative rate of 4.3%. This rate is a conflation of false positives in HG002, false negatives from the sequencing technology and the false negatives from genotyping by LRcaller. Subsampling the file had a minimal effect on the number of variants identified; the sample was reported to be a carrier of 28,919, 28,942, 28,935 and 28,938 for subsampling of 0.1, 0.2, 0.4 and 0.8 times the original coverage, respectively.

We note that our comparisons to HG002 are less accurate compared to when variants are directly ascertained on GRCh38; some variants that can be called in GRCh38 may be missed in GRCh37 and the liftover process may not correctly represent some variants ascertained in GRCh37.

Next, we genotyped the variants discovered in our study within HG002 Tier 1 regions on the HG002 sample. HG002 was found to be a carrier of 10,858 of these variants (as determined by LRcaller). For 10,126 of these variants a corresponding variant was found in the HG002 truth set while for 732 no corresponding variant was found, suggesting a 6.7% false positive rate for our variants within Tier 1 region. A fraction of these may be false negatives in the HG002, but we did not attempt to quantify this fraction. Again, coverage had limited effect on these results, when subsampling HG002 to 0.1 times the original coverage 729 of the variants were not found in the HG002 truth set.

### Phasing and imputation of SVs

For each marker, we produced a total of 13 different genotypes, 3 from GraphTyper/Illumina and 10 from LRcaller/ONT. We phased and imputed all genotyped variants into the haplotypes of 166,281 Icelanders, using a methodology previously described^19,25,26^. We considered variants with an imputation info greater than 0.9 and a leave-one-out R^2^ greater than 0.5 as *imputed*. After imputing our SVs, we also acquired the allele frequencies of the variant alleles, and the reference allele spanning a multi-allelic SV. We use the reference alleles to deduce haplotypes carrying a variant allele of unresolved size or type. Carriers of a variant allele of unresolved type are determined as those not containing the reference allele of a secondary form multi-allelic SV comprising both insertion and deletion variant alleles. We refer to variant alleles with an unresolved size or type as *non-reference alleles*.

### SV Filtering

After genotyping and imputing our merged SV set using both LRS and SRS sequencing we filter them using imputation accuracy and other filters.

i. **Determining the best genotyping for a variant allele:** In order to find the best genotyping for a variant allele, we sort them by their imputation info, and break ties using leave-one-out R^2^, allele frequency, and genotype models, in the given order. We prioritize genotype models as follows: AD, VA, J, VAr, PR for LRS genotyping, and AG, B1 = B2 = B = C for SRS genotyping. Left breakpoint models are preferred over right breakpoint models in LRS genotyping, within a model. Primary form variants from LRS genotyping are given precedence, if imputed. Best genotypes for LRS and SRS genotyping are found separately. Imputed genotypes are preferred, if they exist.
ii. **Assigning quality groups to variant allele genotypes:** We assign genotypes into four quality groups based on their imputation accuracy and allele frequency. Imputed alleles are assigned Group 1 (G1). Alleles with a sufficient imputation information (greater than 0.9) but an insufficient leave-one-out R^2^ (less than 0.5) with allele frequency less than 0.2% are assigned Group 2 (G2). SVs with both an insufficient imputation information and a leave-one-out R^2^ with allele frequency less than 0.1% are assigned Group 3 (G3). The remaining alleles are assigned Group 4 (G4). We created groups 2 and 3 as it is more difficult to phase and impute alleles with few carriers accurately.
iii. **Filtering variant allele quality groups and LRS/SRS genotyping preference:** We accept all G1 alleles into our merged SV set as imputed alleles. For G2, G3 and G4 alleles, we require at least 90% of the individuals they were discovered in to remain as carriers after phasing (see “Phasing and imputation of SVs”). For G4 SVs, we additionally require at least 90% of carriers after phasing to be found among the set of individuals they were discovered in. We assign and filter quality groups in variant alleles independently for LRS and SRS genotyping. If the variant is a TR SV, primary form variants from LRS genotyping are given precedence over secondary form variants from LRS genotyping or SRS genotyping, if imputed. Otherwise we use the same method described in “Determining the best genotyping for a variant allele”. We retained 339,216 SV alleles after this filtering, such that 320,778 were among G1, 4,783 among G2, 6,352 among G3, and 7,303 among G4.
iv. **Linkage Disequilibrium (LD) filtering:** We compute pairwise LD R^2^ measures of variant alleles within 10 kb proximity using their begin sites, in order to remove potential duplicate calls. In order to find alleles in LD, we use an undirected graph *G*(*V,E*) where each υ ∈ *V* vertex represents an SV, and edge *υ* ∈ *E* between vertex *υ*_*i*_ and vertex *υ*_*j*_ is drawn if*υ*_*i*_ and are of the same type, or at least one of them is of unresolved type, and they have an LD R^2^ of 0.8 or greater. Next, we find the connected components (CC) in *G*, and pick one representative for every CC using the same method described in **iii**. We retained 156,687 variant alleles after LD filtering, such that 83,082 and 57,885 are insertions and deletions of specified size, 7,819 and 3,840 are insertions and deletions of unspecified size, and 4,061 are unresolved insertions or deletions.
v. **Filtering agglomerate multi-allelic SV alleles**: We expect the variant alleles of a multi-allelic SV to be found in distinct sets of haplotypes. Due to imperfections in genotyping, this expectation can be violated, resulting in a multi-allelic SV allele with carrier haplotypes of multiple other alleles. We refer to such alleles as *agglomerate alleles*. We seek agglomerate alleles among all SV alleles within 250 bp of each other using their begin sites, which we refer to as *proximal alleles*, for a conservative filtering. To solve this, we compute a Maximal Independent Set (MIS) in graph *G*(*V,E*), where each vertex *υ ∈V* represents a proximal allele, and edge *e* ∈ *E* between vertex *υ*_*i*_ and vertex *υ*_*j*_ is drawn if two proximal alleles have *shared* carrier haplotypes, separately for insertions and deletions. Two alleles have shared carrier haplotypes if

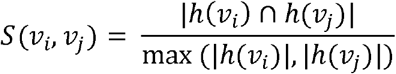

is at least 20%, where *h* (*υ*_*i*_) refers to the set of carrier haplotypes of *υ*_*i*_. As a graph can have many MISs, we use a greedy algorithm where we construct a MIS by iteratively selecting a vertex of minimum degree, and removing it and its neighbors from the graph until the graph it empty^72^. We break ties by prioritizing primary form, and lower frequency alleles, in the given order. Non-reference SV alleles are expected to have shared carrier haplotypes with proximal multi-allelic SV alleles with a to have shared carrier haplotypes with proximal multi-allelic SV alleles with a specified size, therefore they are not included in this step. We perform a separate filtering using all proximal non-reference alleles and use a unresolved type, and they have an LD R^2^ of 0.8 or greater. Next, we find the *S* (*υ*_*i*_)cutoff of at least 80%, to filter near-duplicate non-reference alleles. Using all variant alleles in constructed MISs, we report 133,886 variant alleles, such that 68,595 and 52,075 are insertions and deletions of specified size, 6,455 and 3,574 are insertions and deletions of unspecified size, and 3,187 are unresolved insertions or deletions.

We report the set of variant alleles from our final filtering step as *high-confidence* variants.

### Effect of different overlap parameters in SV merging in number of SVs

We ran our pipeline with SVs discovered and genotyped in chr21 using LRS, which we merged with different distance parameters *D* while finding SV cliques to see their effect in the number of SVs. Initially we ran SV merging with a uniform, i.e. same for within and outside of TRs, *D* of 0.99, 0.75, 0.5, and 0.25, and observed 834, 1,081, 1,584, and 2,269 imputed markers, respectively. Next, we ran SV merging with a *D* of 0.5 for SVs outside of TRs, and 0.25 and 0.15 for SVs within TRs, where we observed 2,273 and 2,554 imputed markers, respectively. We noticed that the increase in the number of imputed markers from using a *D* of 0.5 to 0.25 is explained by the variation within TRs, as shown by the similar number of imputed markers between using a single *D* of 0.25 and dual *D* s of 0.5 and 0.25 (2,269 vs. 2,273). This also showed us that a *D* of 0.5 was indeed over-collapsing distinct variant alleles within TRs. Seeing the increase in imputed markers using dual *D* s of 0.5 and 0.15 (2,273 vs. 2,554) we decided to use this parameter set in our SV merging. We did not pursue further restriction in distance as it can be difficult to reliably genotype markers with very similar sizes in LRS given its error rates.

### Calculating SV counts per individual

We calculate the number of SVs found per individual using a conservative estimate in order not to double count variant alleles. We treat all proximal alleles as a single SV, and count SVs per individuals accordingly.

### Effects of sequencing-related differences to SV numbers in an individual

We assessed the effect of total error rate, aligned coverage, and status of being sheared on both the number of discovered (Fig. S5), and high-confidence variants (see “Calculating SV counts per individual”), per individual (Tables S1, S2). Although all tested variables had a highly significant effect on the number of discovered variants, only aligned coverage had a significant effect on the number of high confidence variants per individual (an increase of 1.83 (0.008%) variants per 1x aligned coverage, p = 0.03).

### Comparison of the merged SV set to other SV datasets

In order to calculate false positive and false negative rates of our high-confidence SV alleles, we developed a statistic, variants inconsistent with HG002 (VIH), and compared our dataset to the SV sets provided by Audano et al.^30^, gnomAD-SV^12^, and Zook et al.^5^ on HG002. We expect most high frequency variants to be present in populations of similar heritage, such as European populations. Variants found in population A but not found in population B can be due to i) fixation of the variant in population B, ii) different representation of the variants between the two datasets, iii) false positive calls in dataset A, and iv) false negative calls in dataset B.

As variant representation differs between variant callers we use a relaxed method for determining whether a variant discovered in dataset A is also present in dataset B, counting the variant as present if there is a variant in dataset B occurring within 500 bp from the start position of the variant discovered in dataset A (given gor files for datasets A and B the command run was: gorpipe “A.gor | join –snpseg –f 500 B.gor”).

i. **VIH statistic – Variants inconsistent with HG002** As HG002 has been extensively characterized within Tier 1 regions we can use it to estimate false positive rates within these regions. We develop the statistic VIH (variants inconsistent with HG002). VIH takes as input two studies, A and B along with their variants and the variant frequencies from study A. VIH assumes that 1) the same classes of variants have been characterized within HG002 as in A and 2) no drift in variant frequency has occurred between A and HG002. We compute

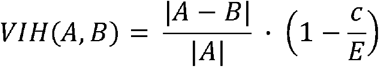

where |.| represents cardinality, |A − B|the number of variants in A missed by study B.*c* is the number of variants in | A − B | found in HG002 and E is the expected number of variants in | A − B | found in HG002 given the variant frequencies in A.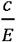 is therefore an estimated true positive rate of the variants in | A − B | and 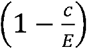 an estimate of false positive rate. We use this statistic as a surrogate for false positive rate for study A, although it may in part be explained by population drift, differences in variant classification between studies and false negatives in the HG002 truth set.
ii. **SVs found by Audano et al. in our dataset** We first compared our study to the one of Audano et al. The study has individuals of diverse ancestry, but we would expect that most variants above 50% frequency to also be present in our study. We find that 93.0% of the variants are present in our study, nominally suggesting a false negative rate of 7.0% for our study. We further classified these variants according to the genomic classification of GiaB for HG002, where Tier1 represents regions where the most reliable calls can be made. As expected the agreement is highest within Tier1 regions, where we find 96.5% of variants to be present within our study, i.e. 3.5% (235) as missing from our study in those regions. 162 of these were found in HG002, compared to 218.2 expected, suggesting that the 3.5% can be partitioned into a 0.9% VIH rate (A = Audano et al., B = our high-confidence SV set), representing population differences or false positive rate for the SV set by Audano et al. and a 2.6% false negative rate for our study, within Tier 1 regions. Similarly, we can find 94.6% of the variants within Tier 1 regions above 10% frequency in the study by Audano et al. This drop can be in part explained by the different ancestry of individuals in the study by Audano et al.
iii. **SVs found by Audano et al. in gnomAD-SV** gnomAD-SV found 58.7% and 62.2% of variants discovered by Audano et al. with frequency greater than 50% and 10%, respectively. When restricting to Tier 1 regions the fraction found in gnomAD-SV was 61.0% and 66.0%, respectively. We computed a VIH rate of 0.0% (A = Audano et al., B = gnomAD-SV) for the SV set by Audano et al. as more variants private to Audano were found in HG002 than expected.
iv. **SVs found by gnomAD-SV in our dataset** We compared gnomAD-SV using variants within Tier 1 regions above 10% frequency to our high-confidence SV set, and observed 9.7% of them to be missing in our study. This rate can be partitioned into a 6.3% false positive rate (using VIH) for gnomAD-SV, and a 3.4% false negative rate for our study using our VIH statistic as described above.
v. **SVs found by gnomAD-SV in Audano et al**. We also compared the same subset of SVs from gnomAD-SV to the study by Audano et al., and estimate a 7.6% false positive rate for gnomAD-SV, using VIH. We believe a false negative rate estimate for the Audano study using this approach is not informative as some of the missing variants can be attributed to the small sample size (N = 15) in the Audano study.
vi. **SVs in our dataset in gnomAD-SV and Audano et al**. Similarly, we compared our high confidence variants within Tier 1 regions above 10% frequency (21,029 variants) to the other studies. 88.7% and 62.7% were found by Audano et al. and gnomAD-SV, respectively. We were not able to use gnomAD-SV to compute VIH (A = our high-confidence SV set, B=gnomAD-SV) in our dataset due to the small fraction of variants found by gnomAD-SV. Of 2,377 of our variants not found in the study by Audano et al., only 331 were found in HG002, compared to 1,195 expected, resulting in a VIH (A = our high-confidence SV set, B = Audano et al.) of 8.2% false positive rate estimate for our dataset within Tier 1 regions. We expect that our method may in some cases call variants that are slightly shorter than 50 base pairs as structural variants. The remaining difference are explained by false positives in our method and possibly some drift in frequency between the Icelandic population and the Ashkenazi population of HG002.

### Distribution of genotype calls in parent-offspring trios

We computed the distribution of SV genotype calls in all parent-offspring trios included in the study (Table S3) as quality metric for our high confidence set. The Mendelian laws of inheritance dictate that both alleles have equal probability of transmitting and therefore can predict the distribution of offspring given the parent genotypes. We use 0 to denote the reference allele and 1 to denote the alternative allele. For example, if both parents are called as heterozygous (0/1) then we expect the offspring genotypes to be homozygous reference (0/0), heterozygous (0/1), and homozygous SV allele (1/1) in 25%, 50%, and 25% of the time, respectively. Distributions closer to the expected probabilities indicate more accurate genotype calls.

### Polymerase Chain Reaction (PCR) verification of SVs

We randomly selected 80 SVs (10 for each SV category we defined) among our imputed SV set for PCR and agarose gel electrophoresis. We defined our SV categories with respect to their underlying event (insertion or deletion), length (less than or greater than 1 kb), and whether they are within or outside a TR region, amounting to 8 categories in total. All non-carriers, heterozygous carriers, and homozygous carriers were randomly selected from our imputation set of 166,281 individuals.

A single primer pair was designed flanking each small SV (less than 1 kb in length) using the Primer3 program. For large SVs (greater than 1 kb in length), two pairs were designed for co-amplification such that the first pair would amplify only carrier chromosomes and the second would amplify only non-carriers chromosomes. In both scenarios the subsequent PCR resulted in different sizes of the amplified fragments, depending on the genotypes of the samples. PCR and agarose gel electrophoresis were carried out using standard protocols on DNA from two individuals of each expected genotype: non-carriers, heterozygotes and homozygotes (when available). The fragment sizes were estimated by running the amplicons on a gel adjacent to a DNA ladder (FastRulerTM Middle Range DNA Ladder, ThermoFisher Scientific).

Of the 80 SVs, 10 failed in PCR primer design. Among the remaining 70 SVs for which PCR verification was attempted, 7 failed to give a PCR product of the predicted size, 60 were confirmed to contain a SV at the predicted site, and 3 only showed the expected non-carrier bands. 4 of the tested SVs have an updated start site in our current dataset (Table S4, Supplementary Data S3).

### Characterization of insertions

We classified insertions into three groups; tandem duplications (TD), retrotransposable elements (RE), and other insertions (INS) as follows. We mapped all insertion sequences to GRCh38 using minimap2 and calculated the distance between their mapping locations and original begin sites where they were discovered. All insertions with this distance less than 2 times the insertion length, and with a query coverage of 80% are accepted as TD. We mapped the remaining insertion sequences to human transcripts using refseq annotations^77^ and accepted insertions with a query coverage of 80% to a single transcript as RE. The remaining insertions are annotated as INS.

### Analysis of genomic regions

Epigenome data from the NIH Roadmap Epigenome Mapping Consortium^73^ and Encode project^74^ for 11 histone marks analysed by ChIP-seq, together with open chromatin regions analysed by DNase-seq, have been integrated into 25 discrete chromatin states through use of the ChromHMM software^75^. The integrated data (referred to as the ChromHMM 25-state model) were retrieved from the official Roadmap Epigenomics website (www.roadmapepigenomics.org). ChromHMM annotations for primary tissues and cells (excluding those for immortalized cell-lines) reflect the union for each of the following chromatin states: Bivalent (PromBiv), Active and Weak enhancers (enhancers class A1 and A2, W1, W2), Transcription start sites and promoters (TssA, PromU, PromD1, PromD2), Poised (PromP), PRC2 (polycomb group repressor complex 2 marked regions), Heterochromatin (Het) along with Transcribed regions (Tx).

Double strand break regions (DSBs) determined from DMC1 binding in spermatocytes were described by Pratto et al.^31^, liftover from GRCh37 to GRCh38 was applied before analysis. Recombination hotspots are from Halldorsson et al.^76^. Coding exons were determined using refseq annotations^77^. Cpg islands were taken from the UCSC browser^78^. For each genomic attribute overrepresentation is computed as (A/B)/G, where A is the number of SVs with a start point within regions annotated with the attribute in question, B is genomic size of those regions and G is the genomic average number of SVs starting at a given position.

For each genomic attribute the fraction of rare variants (FRV) is computed as (A/B)/F, where A is the number of rare SVs (frequency below 1%) that overlap the region of interest, B is the number of SVs that overlap the genomic attribute and F is of the fraction of SVs that are rare in the complete dataset.

### Confidence intervals and p-values

All confidence intervals are 95% confidence intervals. Confidence intervals and p-values are computed by bootstrapping 1,000 sets of the same size as the original statistic is computed over, sampled with replacement from the set of SVs. The statistic in question is computed within each set, creating a list of 1,000 statistics. Following the sorting of this list, the lower bound of the confidence interval is computed as the mean of entries 25 and 26 and the upper bound is computed as the mean of entries 975 and 976. Bootstrapping is also used to compute p-values, the number of times an event as or more extreme than the event observed is tabulated and multiplied by 2 to represent two sided statistic. In the case when as extreme event was not observed the p-value is reported as < 0.002.

### Association testing

We tested our SVs for association with LDL levels based on the linear mixed model implemented in BOLT-LMM^79^. We used BOLT-LMM to calculate leave-one-chromosome out (LOCO) residuals which we then tested for association using simple linear regression. A generalized form of linear regression was used to test for association of phenotypes with SVs. We assume that the phenotypes follow a normal distribution with a mean that depends linearly on the expected allele at the variant and a variance-covariance matrix proportional to the kinship matrix^80^. We used linkage disequilibrium (LD) score regression to account for distribution inflation in the dataset due to cryptic relatedness and population stratification^81^. The inflation factors were computed from a set of SNP and indel sequence variants. Using a set of about 1.1 million SNP and indel sequence variants, we regressed the χ2 statistics from a genome-wide association scan against LD score and used the intercept as correction factor. Effect sizes based on the LOCO residuals are shrunk and we rescaled them based on the shrinkage of the 1.1 million variants used in the LD score regression.

### Comparison to GWAS catalog

We downloaded version 1.0 of the GWAS catalog with all associations (see URLs) on July 23^th^ 2020 (`gwas_catalog_v1.0-associations_e100_r2020-07-14.tsv`). SNPs and indels in the GWAS catalog were matched with in-house SNPs using exact coordinate matching and two markers were assumed to be the same if they had the exact same coordinate in GRCh38.

An inhouse tool was used to compute correlations between SNPs and indels imputed into 166,281 Icelanders and SVs imputed into the same set. Correlations were limited to windows of 500 kb, such that a correlation between a SNP/indel and a SV is observed if and only if they are within 500 kb of each other. We provide our results in Supplementary Data S4.

### Calculating the geographic distribution of the carriers of the *PCSK9* deletion

In the last century the percentage of the Icelandic populations living in the capital area has gone from 22.7% (21,441 out of 94,436 in 1920) to 62.4% (222,779 out of 356,991 in 2019). The extent of these migrations suggest that the geographic distribution of the variant might have changed from the historical geographic distribution in the last century. In order to obtain geographic distributions closer to the historical geographic distribution, for each carrier and non-carrier we inferred the carrier status of their grandparents and calculated allele frequencies in regions based on the grandparents. The two haplotypes of every chip typed individual had been assigned a parent of origin and we estimated the allele frequency within the recorded birth-county of each grandparent by assuming there was a 1/2 chance that parental allele came from that grandparent. We grouped the 22 Icelandic counties into 11 regions as presented in Table S5.

### Association testing for the number of motifs in *ACAN*

Using the SV alleles we identified in an ACAN VNTR (Table 1), we calculated expected motif change per haplotype in an individual as

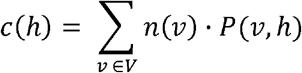

where *υ* ∈ *V* represents the SV alleles, *n*(*υ*) the number of motif change in SV allele, *υ* (negative for deletions). *P*(*υ,h*) is haplotype carrier probability for haplotype *h* carrying the which we calculate by rounding the allele lengths divided by the motif length to an integer SV allele *υ*, calculated during imputation (see “Phasing and imputation of SVs”). We performed a unit-based normalization of the *c*(*h*) values to use as haplotype carrier probabilities in association testing.

### URLs

SViper, original repository: https://github.com/smehringer/SViper

Scrappie, original repository: https://github.com/nanoporetech/scrappie

GWAS catalog: https://www.ebi.ac.uk/gwas/docs/file-downloads

Chemagen: https://chemagen.com

HG002 truth set: https://ftp-trace.ncbi.nlm.nih.gov/giab/ftp/data/AshkenazimTrio/analysis/NIST_SVs_Integration_v0.6/HG002_SVs_Tier1_v0.6.vcf.gz

HG002 ONT dataset: https://community.nanoporetech.com/knowledge/datasets/hg002

## Supplementary Material

**Figure S1:**
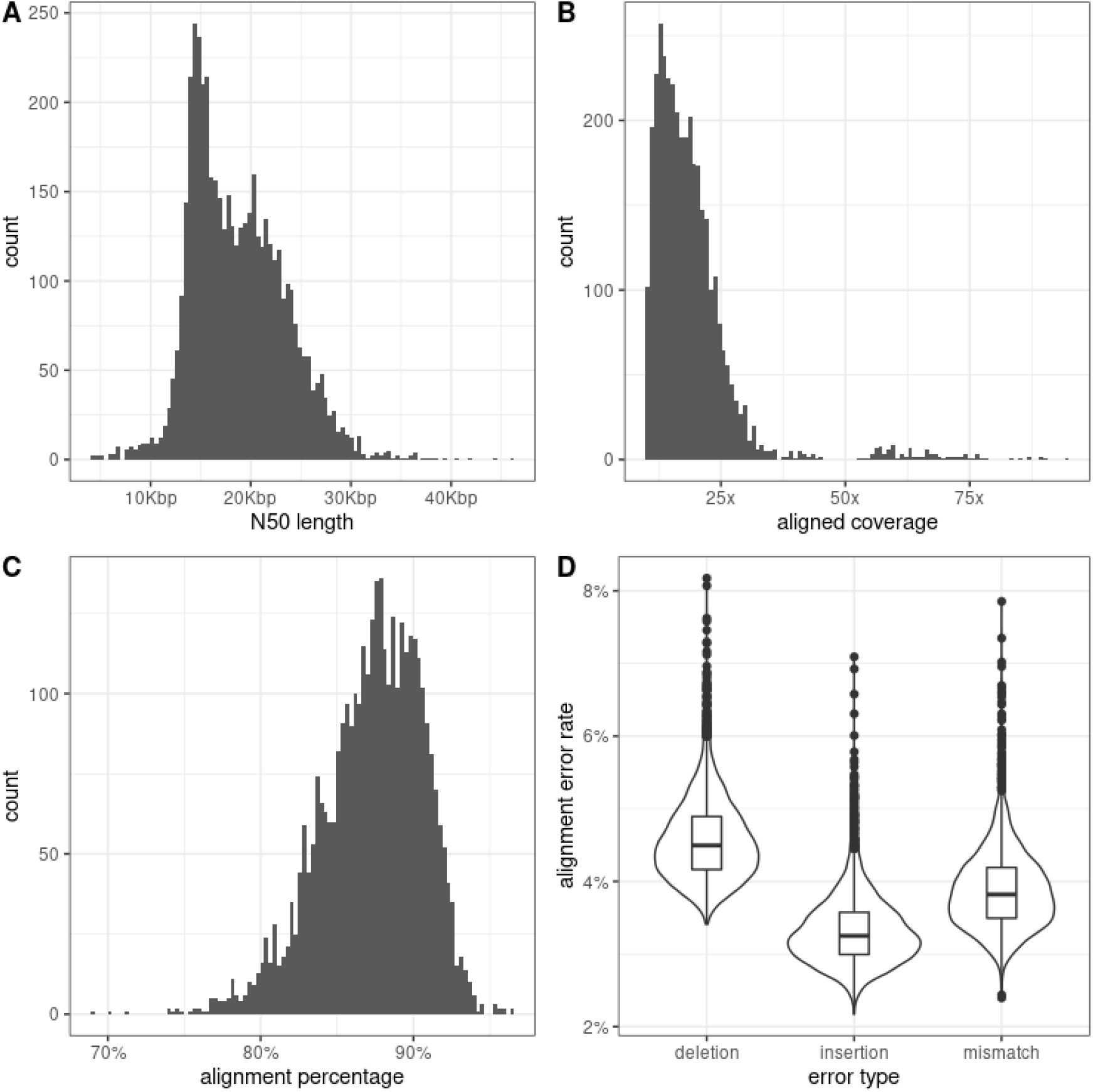
Oxford Nanopore Technologies (ONT) long-read sequencing statistics. (A) N50 length per flowcell (n = 4,757 flowcells) prior to GRCh38 alignment. (B, C, D) Aligned coverage, alignment percentage, and error rates stratified by type, per individual (n = 3,622 individuals). Statistics are computed over sequenced reads longer than 3000 bp.

**Figure S2:**
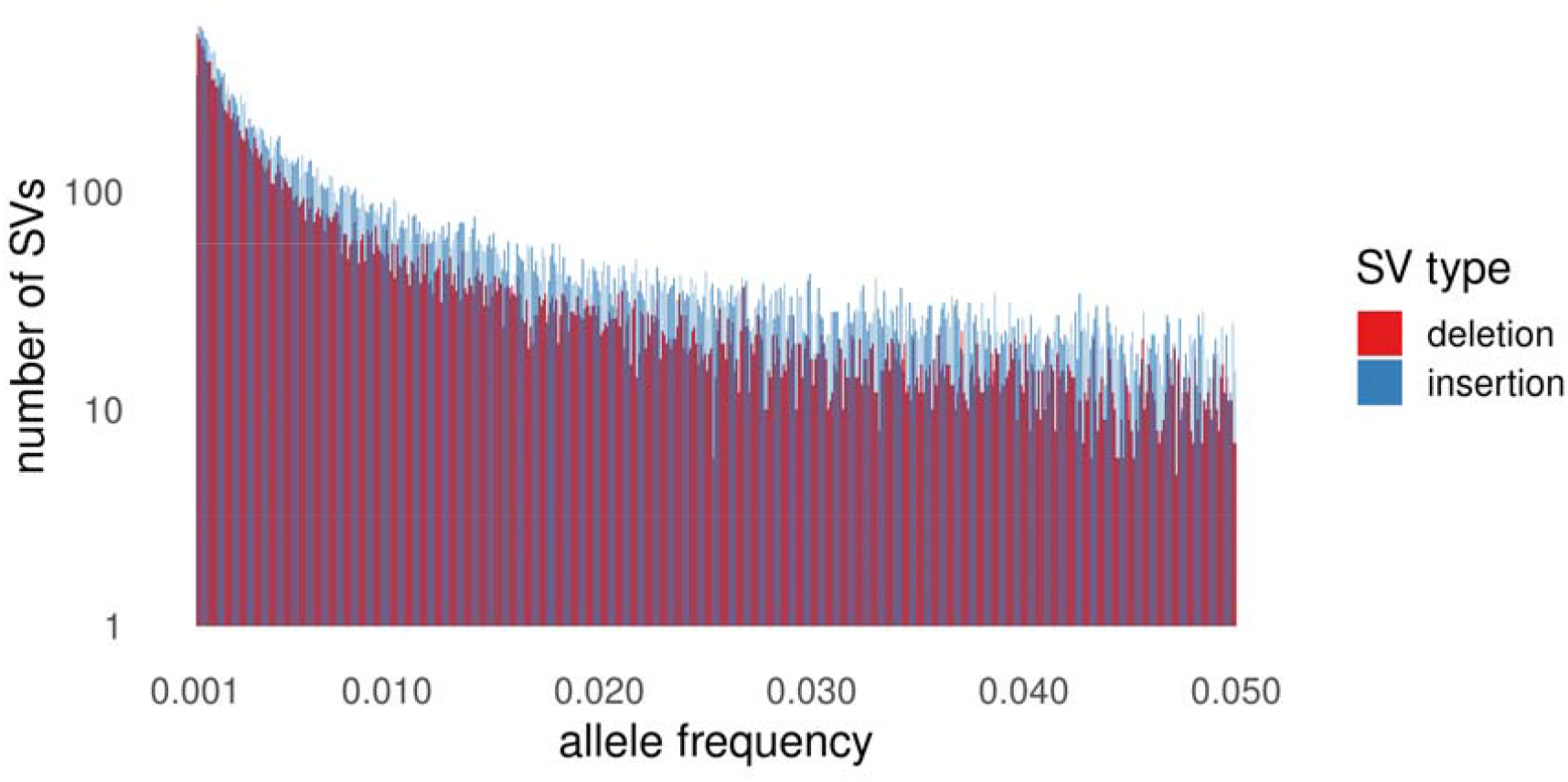
Allele frequency distribution of SVs binned at 0.01% for alleles with 0.1% to 5% frequency.

**Figure S3:**
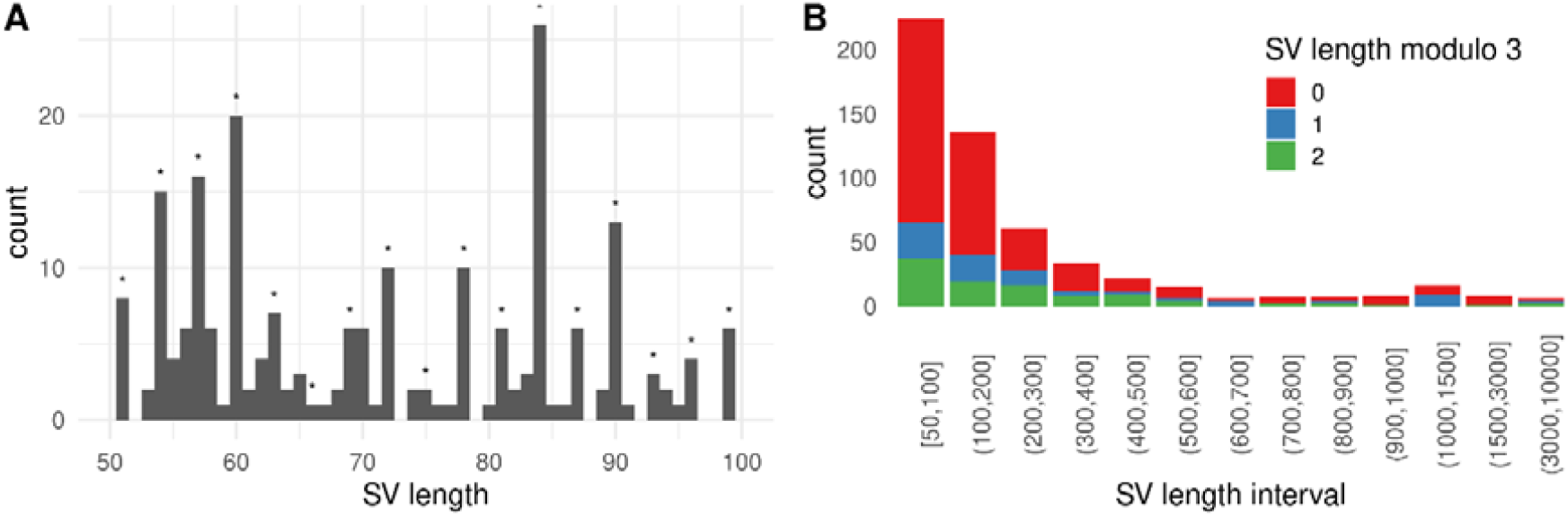
Length and modulo distributions of structural variants (SVs) that are contained within exons. (A) Length distribution of SVs with lengths between 50 and 100. Stars denote lengths divisible by 3. (n = 224 markers) (B) Modulo distribution of SV lengths across length intervals. (n = 549).

**Figure S4:**
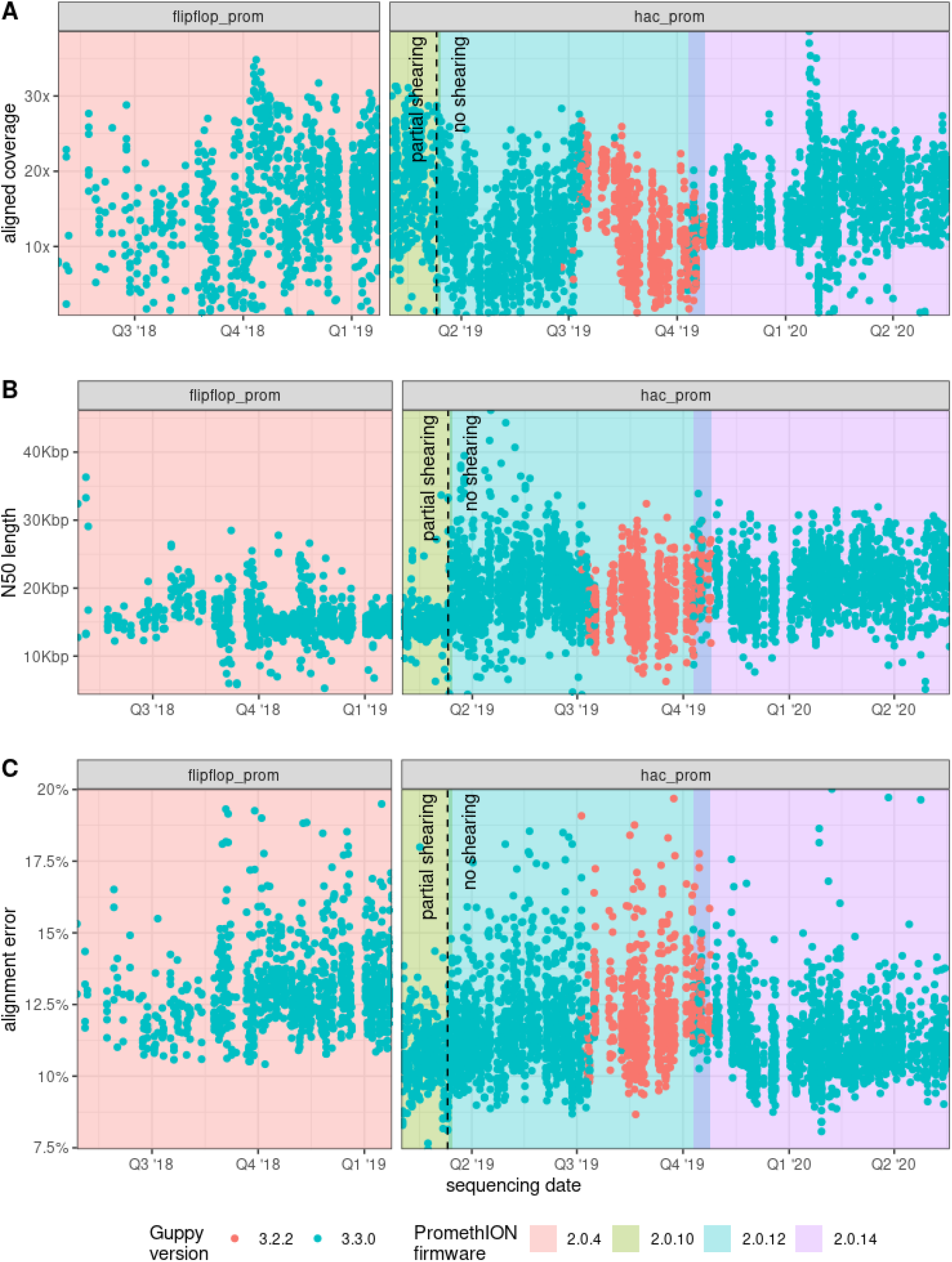
A total of 4,757 flowcells were analyzed for 3,622 individuals from May 2018 until May 2020. In March 2019 we stopped partially shearing the DNA, resulting in an increased N50. For PromethION flowcells with firmware 2.0.4 we used the *flipflop* basecalling model, and the *hac* basecalling model for firmware 2.0.10 and later.

**Figure S5:**
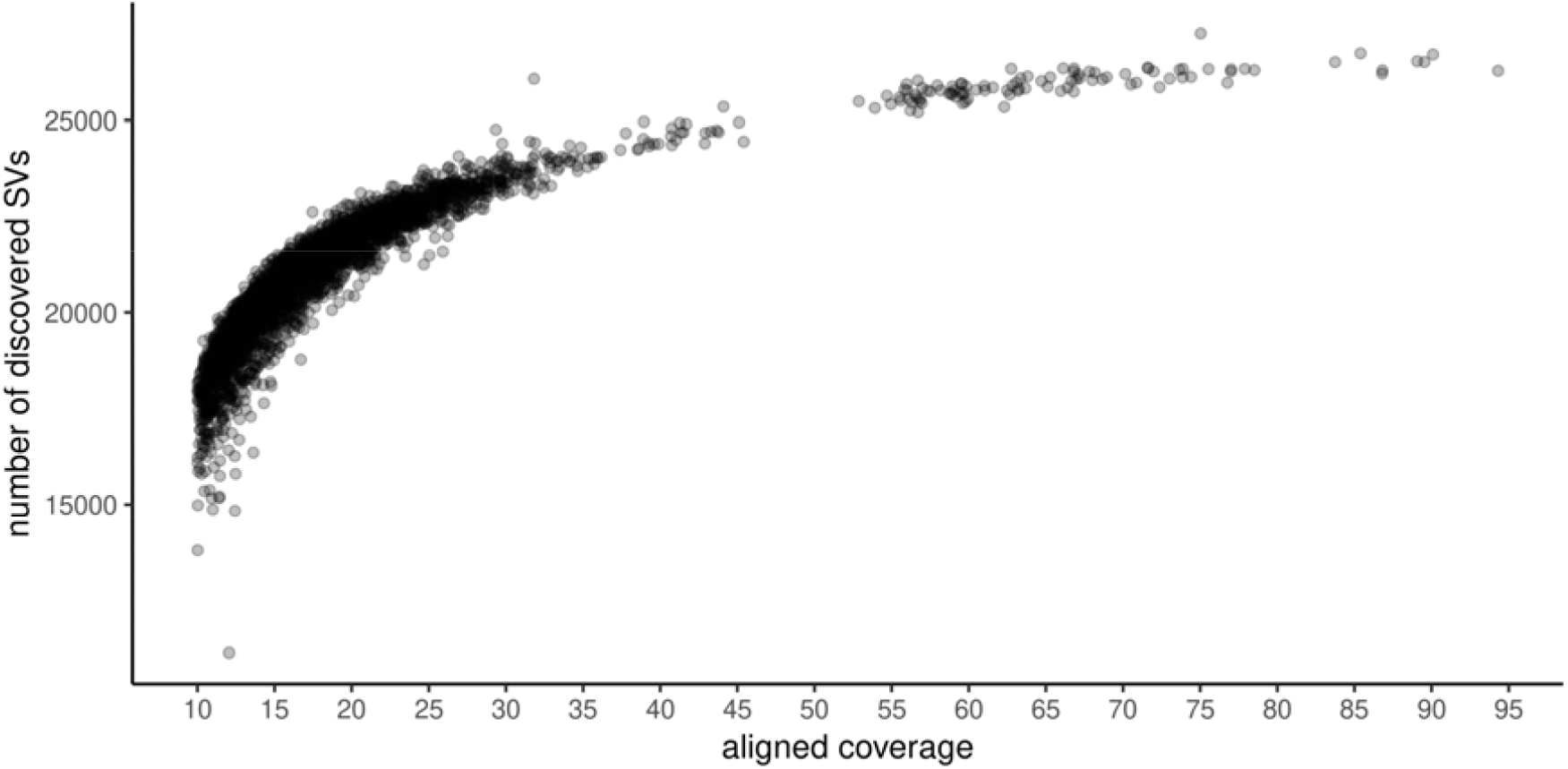
Number of discovered structural variants (SVs) vs. aligned coverage, per individual (N = 3,622 individuals).

**Figure S6:**
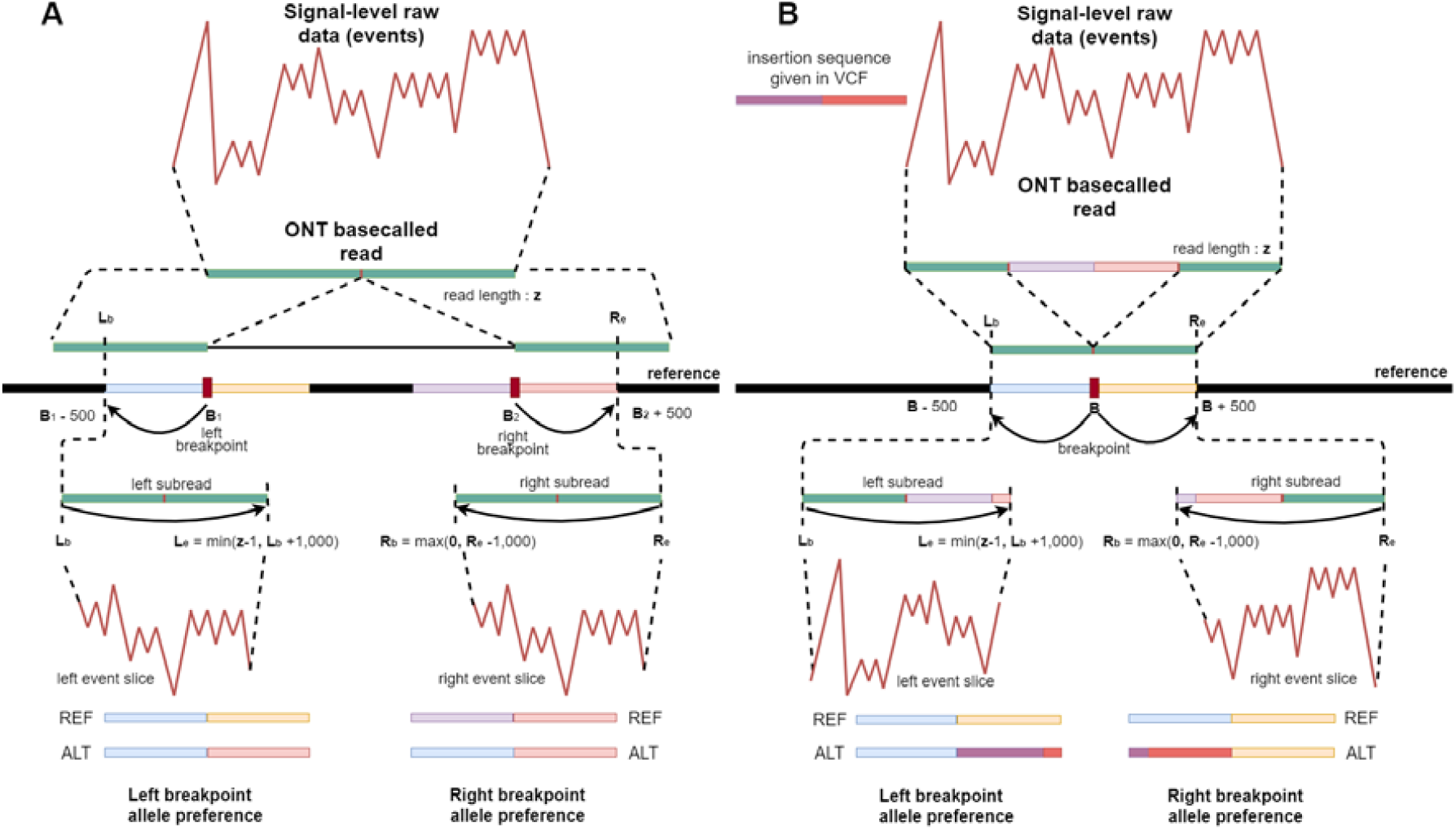
SquiggleSVFilter overview. Given a candidate structural variant (SV), and an SV supporting read, SquiggleSVFilter first identifies the subread of the ONT basecalled read overlapping the SV, using the reference alignment BAM file. Next it finds the squiggle slice of the identified subsequence using the event table. For both the left and right flanks around the variant, it determines the reference and alternative sequences given the candidate variant, and computes their raw data-vs-sequence log likelihood scores with the squiggle slice. A sufficiently high log likelihood score difference for the alternate allele marks the read as an SV supporting read.

**Figure S7:**
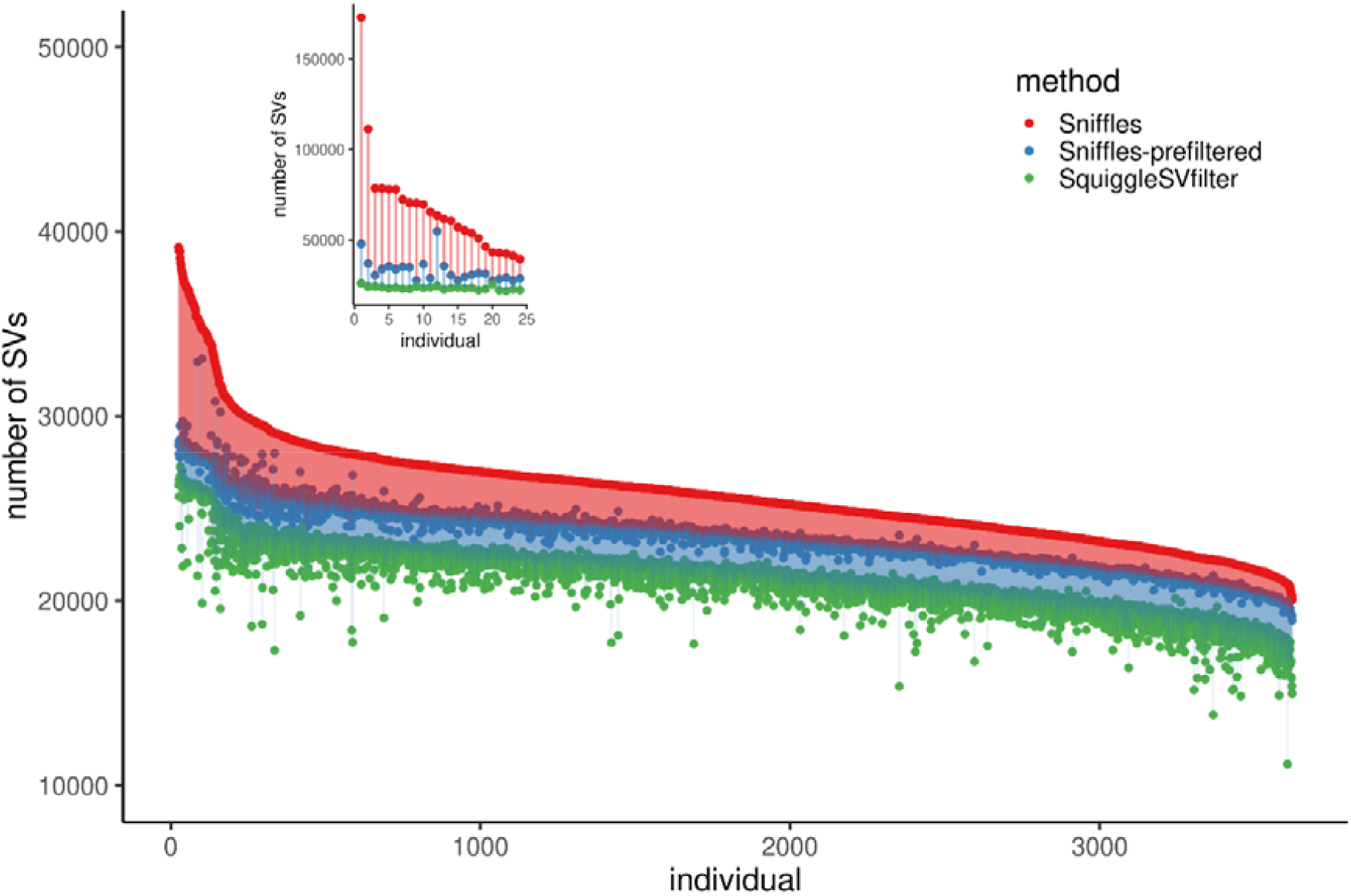
Structural variant (SV) filtering using SquiggleSVFilter. Per indvidual SV calls are shown for Sniffles (red), a local alternative allele ratio based pre-filtering (blue), and SquiggleSVFilter (green), where each subsequent step uses the output of the previous step as input SV set.

**Figure S8:**
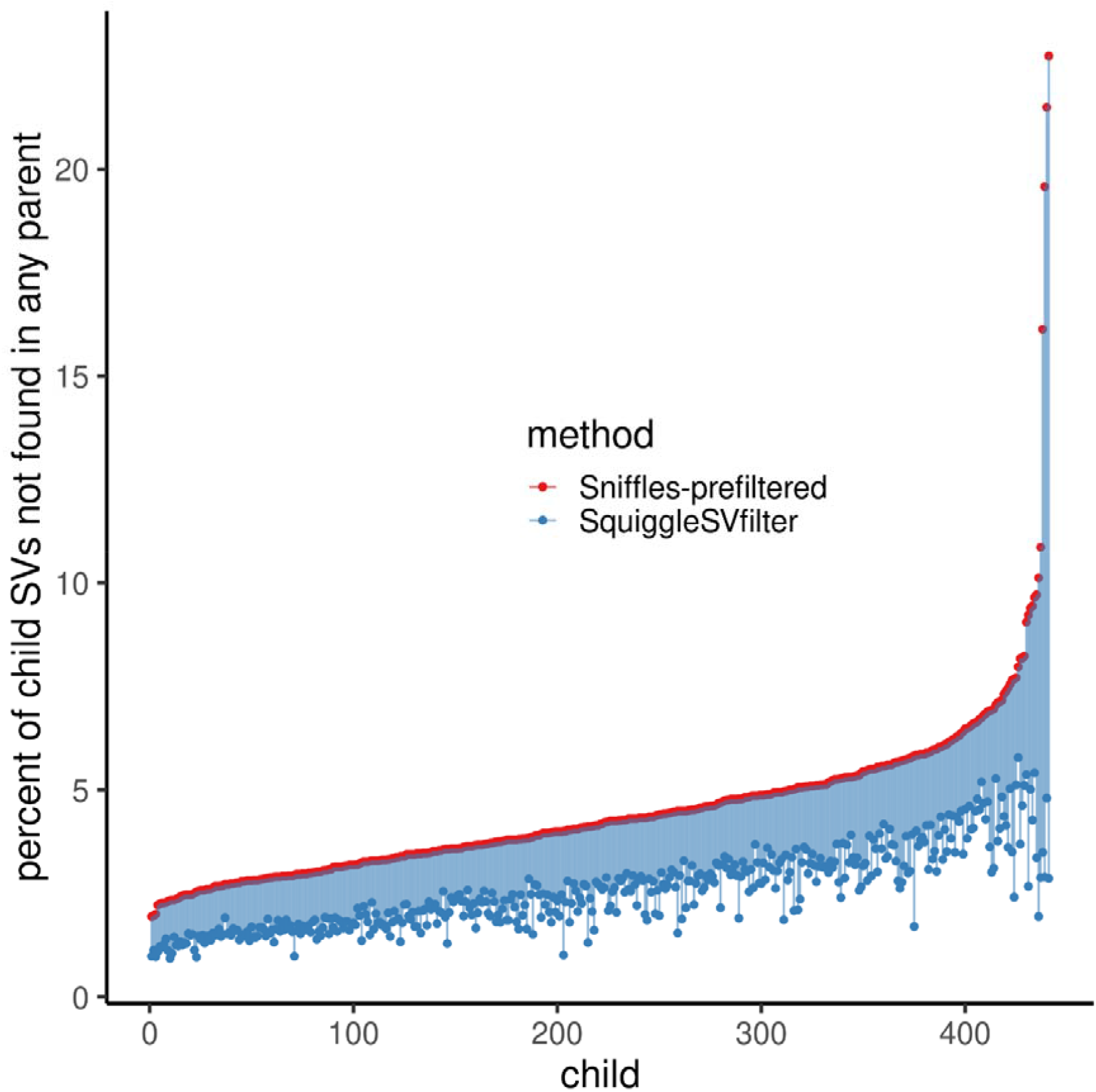
Percent of child SVs not found in any parent in 441 children on the prefiltered Sniffles (red), and on SquiggleSVFilter filtered (blue) calls.

**Table S1:** Effect of aligned coverage, total error rate, and unshearedness to number of discovered SVs per individual (N = 3,622 individuals).

| Term | estimate | pvalue (negative log) |
| --- | --- | --- |
| aligned coverage | 149 | 2098.0 |
| total error rate | -343 | 345.3 |
| unshearedness | -821 | 235 |

**Table S2:** Effect of aligned coverage, total error rate, and unshearedness to number of high-confidence SVs per individual (N = 3,210 individuals, 412 individuals not genotyped using short read sequencing (SRS) data ommited).

| Term | estimate | pvalue |
| --- | --- | --- |
| aligned coverage | 1.83 | 0.03 |
| total error rate | 0.317 | 0.96 |
| unshearedness | 4.09 | 0.82 |

**Table S3:** Distribution of structural variant (SV) genotypes in parent-offspring trios, in percentages. Rows and columns denote parent and offspring genotypes, respectively. We use 0 to denote the reference allele and 1 to denote the SV allele.

|  | 0/0 | 0/1 | 1/1 |
| --- | --- | --- | --- |
| 0/0+0/0 | 99.17% | 0.80% | 0.02% |
| 0/0+0/1 | 52.14% | 46.71% | 1.15% |
| 0/0+1/1 | 8.89% | 85.24% | 5.87% |
| 0/1+0/1 | 24.00% | 54.72% | 21.27% |
| 0/1+1/1 | 2.67% | 49.24% | 48.09% |
| 1/1+1/1 | 0.42% | 3.81% | 95.77% |

**Table S4:** Updated SV start sites for 4 SVs used in PCR verification.

| Sites used in PCR verification | Sites in current dataset |
| --- | --- |
| chr11:103330538 | chr11:103330468 |
| chr16:90023595 | chr16:90023796 |
| chr17:73758029 | chr17:73757958 |
| chr2:238637907 | chr2:238637955 |

**Table S5:** Geographic distribution of the carriers of the PCSK9 deletion across 11 regions from 22 Icelandic counties.

| County | Carrier chromosomes | N chromosomes | Fraction |
| --- | --- | --- | --- |
| Unknown | 3 | 12256 | $2.4 \cdot 10^{-4}$ |
| Gullbringusýsla | 59.5 | 98652 | $6.0 \cdot 10^{-4}$ |
| Kjósarsýsla, Borgarfjarðarsýsla, Mýrarsýsla | 14 | 15358 | $9.1 \cdot 10^{-4}$ |
| Snæfellsnessýsla, Dalasýsla | 0.5 | 19117 | $2.6 \cdot 10^{-5}$ |
| Barðastrandasýsla, V-Ísafjarðarsýsla, N-Ísafjarðarsýsla, Strandasýsla | 3.5 | 37719.5 | $9.3 \cdot 10^{-5}$ |
| Húnavatnssýsla, Skagafjarðarsýsla | 15 | 23652 | $6.3 \cdot 10^{-4}$ |
| Eyjafjarðarsýsla | 2 | 28754.5 | $7.0 \cdot 10^{-5}$ |
| N-Þingeyjarsýsla, S-Þingeyjarsýsla | 0 | 16433 | 0 |
| N-Múlasýsla, S-Múlasýsla | 5 | 25614 | $2.0 \cdot 10^{-4}$ |
| A-Skaftafellssýsla, V-Skaftafellssýsla | 0 | 9843 | 0 |
| Rangárvallasýsla | 9 | 19628 | $4.6 \cdot 10^{-4}$ |
| Árnessýsla | 3.5 | 17014 | $2.1 \cdot 10^{-4}$ |

## Supplementary Data

Supplementary Data S1: Sequencing-related information of 4,757 flowcells from 3,622 individuals.

Supplementary Data S2: Summary level data of high-confidence SVs. Supplementary Data S3: Primer sequences and results from PCR validation.

Supplementary Data S4: 5,238 SVs in strong linkage disequilibrium (LD) with GWAS catalog variants and related data.

Supplementary Data S5: List of genes with at least 1 homozygous carrier of a rare high impact SV allele in our study.

Supplementary Data S6: VCF file (and index) for the high-confidence SV alleles.

Supplementary data will be available upon the conclusion of peer-review and the publication of the manuscript.

